# From Recoverability to Fidelity: Structured Visual Evidence Modeling for EEG Image Reconstruction

**DOI:** 10.64898/2026.11.25.769109

**Authors:** Songjun Du, Bin Tang, Guifeng Deng, Shuying Rao, Junyi Xie, Xi’ang Chen, Haiteng Jiang

## Abstract

A realistic EEG reconstruction can depict the right concept while missing the viewed instance. We introduce Recovery-Based Visual Evidence Modeling to test three aspects of within-concept reconstruction: recovery of spatial detail, reliability of fixed instance decisions, and preservation of instance evidence during generation. In THINGS-EEG2, we refit short-window readouts with one physical session held out at a time. For three participants, post-onset spatial detail outperforms both same-concept exchange and coarse interpolation, and five of six paired post-minus-pre gains have simultaneous intervals above zero. External participant and dataset tests do not establish generalization. On a controlled generation panel for one participant, rescoring with a 128 128 DAv2 joint-assignment endpoint reduces the Own-minus-Exchange advantage by 24.83 percentage points from input to output. Independent state choices remain inconclusive, and visibility in natural outputs is unmeasured. For fixed short-window spatial decisions from held-out sessions, signed semantic support retains 54.46% accuracy at half coverage, versus 52.44% for hash selection and 55.22% for spatial plane margin. The gain over hash survives adjustment; advantages over plane margin or binary agreement are unestablished. These experiments show where a specific readout supports instance detail and where evidence for downstream fidelity and observation-specific confidence remains limited.

## 1. Introduction

An EEG reconstruction can show a dog while changing its coat, pose and surroundings (Fig. 1). To test which instance the recording supports, we compare the spatial detail read from EEG with two same-category references and repeat that comparison after generation. Recent EEG-to-image methods align non-invasive neural recordings with pretrained visual representations and diffusion priors to synthesize plausible scenes [18, 30]. Their realism leaves a fundamental question unanswered: Which details in a reconstruction are supported by the current neural recording, and which are supplied by the generator’s prior? Image-restoration theory separates perceptual realism from reference fidelity [1], and brain-reconstruction studies show why compelling outputs require careful tests of stimulus correspondence [23].

**Figure 1.**
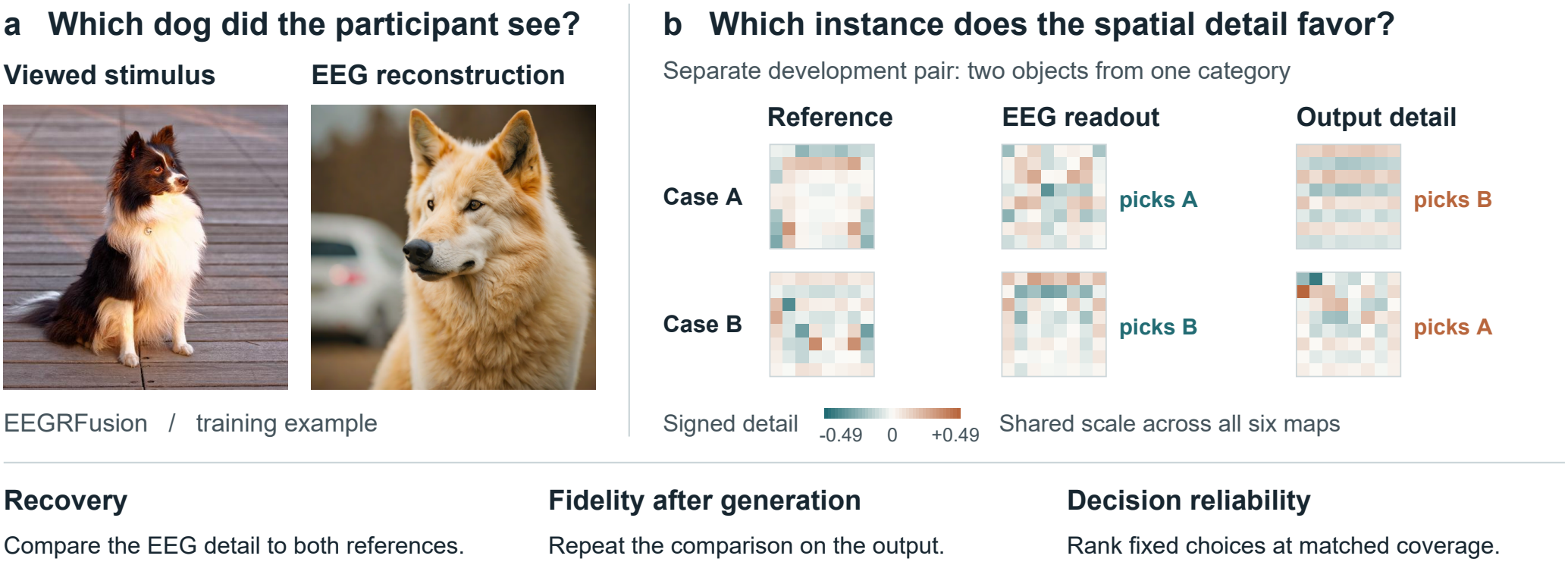
From a recognizable category to a test of instance fidelity. (a) A seated dog becomes a close-up of a light-coated dog in a selected EEGRFusion training example from THINGS-EEG2 [9, 12]. (b) In a separate development pair, each row shows the same 8 8 spatial detail (*R*_8_) measured in the stimulus, read from EEG and measured in the generated image. The EEG readout favors the viewed reference in both rows. Scoring the generated images with the archived DA3 measurements reverses both choices. These cases illustrate the comparison. Fig. 2 defines decision reliability, and Fig. 5 reports the aggregate generation test.

Within-concept instance discrimination makes this question testable by holding the shared concept constant. The challenge is to determine whether a particular layout or spatial detail is grounded in the current recording, and whether generation preserves that distinction. Our premise is that instance evidence is relative to the visual difference being tested: equal recovery errors can have different consequences depending on their direction between candidate images. Individual visual representations differ [2], and prestimulus activity modulates human visual EEG responses [13], motivating measurement under declared participant and observation conditions. Following recovery-versus-use analysis [15], we separate evidence accessible to a readout from evidence realized downstream.

We present Recovery-Based Visual Evidence Modeling, an empirical diagnostic framework connecting multi-scale attributes, same-concept instance distinctions and decision reliability. Building on fine-grained EEG evaluation [27, 31], we follow the same visual distinction through recovery, generation and reliability assessment. The experiments yield three findings.

### 1. Short-window readouts recover instance detail under session holdout

We separate coarse depth layout (*D*_4_) from residual detail (*R*_8_). In three THINGS-EEG2 participants [9], 0–160 ms readouts fitted on other sessions exceed both same-concept exchange and coarse interpolation. We compare the same events before and after onset and report the external protocols where recovery remains unresolved.

### 2. Joint depth evidence weakens along the tested generation pathway

At fixed recipient semantics, layout and noise, we compare the spatial correspondence of Own and exchanged conditions before and after generation. The joint DAv2 effect persists at finer scoring grids that respond to known flips. Independent state decisions and an equal-dose visual-input control limit the interpretation of this effect.

### 3. Held-out reliability gains depend on the comparator

On fixed sub-08 short-window decisions, signed semantic support outperforms hash selection at the same retention but has no established advantage over plain spatial margin or binary agreement. Directional covariance has no established gain over isotropic error. Same-image donor controls also leave observation-specific gains unresolved.

The benchmark separates three questions that visual quality alone cannot resolve: what EEG supports, which instance decisions can be trusted, and what a generator preserves. By linking these questions through explicit visual distinctions and controlled comparisons, it makes failures of instance fidelity measurable and provides a basis for testing more faithful neural reconstruction systems.

## 2. Related Work

### Fine-grained visual EEG decoding

ATM and CognitionCapturer combine EEG alignment with visual representations and generative priors [18, 30]. Hierarchical representations and structural conditions provide additional routes to visual reconstruction [21, 32]. AttrEcho’s preprint studies explicit attributes through within-concept comparisons, controlled representational analysis and complementary encoding; EEG-EditBench’s preprint probes controlled visual alternatives [27, 31]. Our framework connects these visual distinctions to observation-level risk and reconstruction fidelity.

### Recovery and downstream use

Kamath et al. recover content from frozen text representations and compare recovery with controlled multimodal matching [15]. Their separation of retained content and downstream use motivates our EEG analysis of stimulus attributes. Probe capacity also affects what a recovery score reveals, motivating control tasks [11]. Representational modeling relates encoding and similarity analysis through second-moment geometry [4]; noise-unbiased distances require appropriate independent partitions [5]. We specify participant, recording and readout conditions when interpreting attribute recovery.

### Individuality and confidence

Individual visual geometry varies in fMRI [2]; averaging and classifier choices affect M/EEG decoding [10]. Input-dependent uncertainty has also been studied for depth and segmentation [16]. ConfidNet learns confidence from features, and TrustScore assesses agreement with training-data neighborhoods [3, 14]. Selective prediction connects rejection to error at retained coverage [8]. We test the predictive value of attribute-risk supervision against full-representation quality heads and ordinary low-dimensional alternatives.

### Reconstruction fidelity

LPIPS and DreamSim compare images at perceptual levels [7, 33]. BASIC’s preprint evaluates stimulus correspondence at multiple structural and semantic granularities [26]. These measures evaluate an image after generation. Our prospective track predicts a declared output criterion before generation. Candidate-known confidence and gallery-free EEG prediction have different input permissions and are reported separately.

## 3. Recovery-Based Visual Evidence Modeling

### Data and representations

For participant *s* viewing image *I*_*i*_, *X*_*s,i,r*_ is presentation *r* after inherited preprocessing and training-fitted channel normalization [9]. Its processed input has 63 channels and 250 samples at 250 Hz, spanning 0–0.996 s after onset. Four presentations of the same image are averaged before encoding:

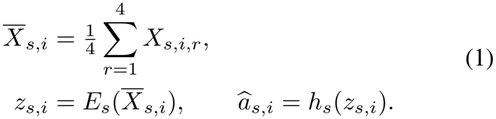

Frozen ATM features *z*_*s,i*_ have 1,440 dimensions. Only *h*_*s*_ is fitted: a linear head or a 256-hidden-unit MLP predicting 336 coordinates, with 16, 64 and 256 entries for *D*_4_, *R*_8_ and *R*_16_. The original 1,024-dimensional semantic projection remains separate; *f*_*s,k*_ = *h*_*s,k*_ º *E*_*s*_ denotes the composite mapping.

### Image roles and instance queries

Stimulus attributes *a*_*i,k*_ = Φ_*k*_(*I*_*i*_) supply supervision and references. A directed query compares one four-presentation EEG readout against two same-concept references; true identity is used only for correctness evaluation. Additional candidates reuse that observation. We trace spatial conditions through generation and assess reliability (Fig. 2). The session-held-out analyses use the short-window protocol described below; the generation section also summarizes a separate EEG-only fidelity prediction study. The Supplementary Material details participant-specific preprocessing and presentation eligibility.

**Figure 2.**
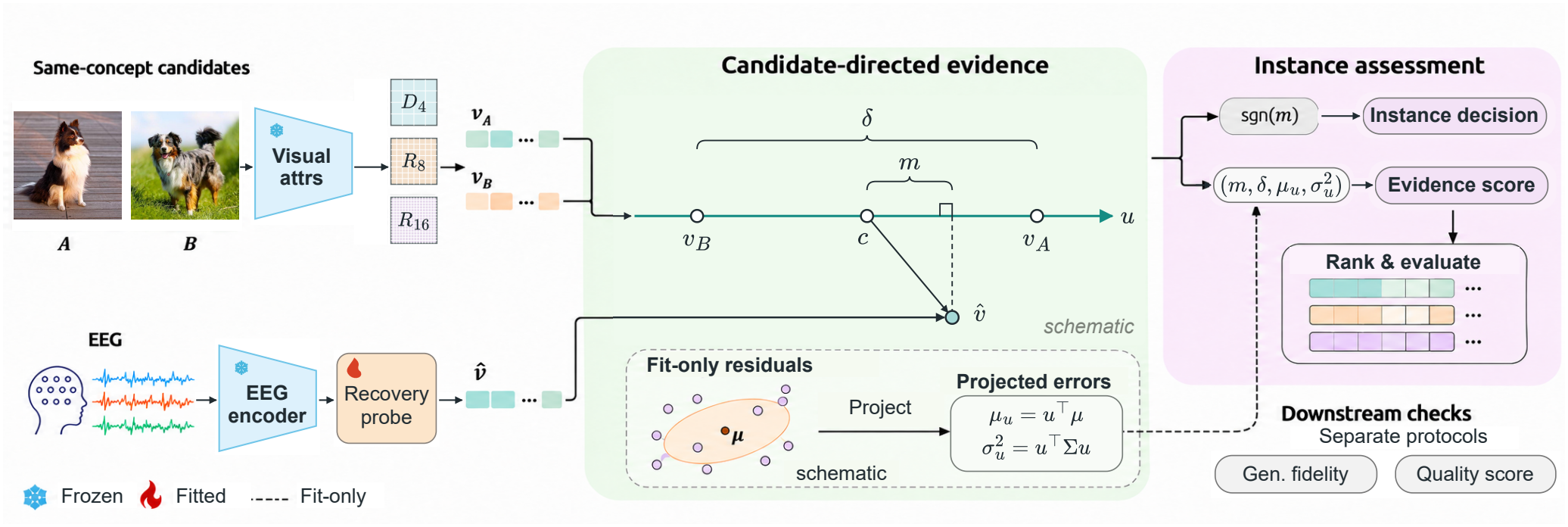
Candidate-directed visual instance evidence. Same-concept candidate attributes define direction *u* and midpoint *c*; EEG recovery gives margin *m*. Fitting-only residuals provide projected bias *µ*_*u*_ and variance 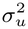 for ranking the fixed instance decision. Recovery, ranking and downstream realization are evaluated separately. Geometry and residuals are schematic; candidate thumbnails are illustrative THINGS-EEG2 training stimuli [9].

### 3.1. Explicit visual coordinates

Explicit feature spaces specify which stimulus properties a neural readout tests [22]. Spatial coordinates make that distinction inspectable. Let *D*_16_(*I*) be the fixed teacher’s normalized 16 *×* 16 relative-depth field, *P*_*n*_ be block-average pooling to *n × n*, and *U* be nearest-neighbor expansion by two. We define

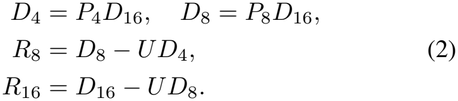

The residuals isolate detail beyond the coarser parent in this representation. We apply identical operators to predictions and references. Existing spatial decoders predict a full 16 *×* 16 field; the measurement decomposition does not change that output into an 8 *×* 8 architecture. A fixed visual embedding provides a complementary semantic attribute.

Spatial targets are standardized using probe-fitting images only. The separate EEG-only fidelity evaluator uses the frozen pre-prior semantic representation. The original generation interface uses Depth Anything V2-Small [28], while new probes use fixed cached Depth Anything 3 attributes [20]. Historical DA3 output scoring used uncontrolled teacher randomness. The seeded replay tests measurement repeatability on fixed cached RGB; DAv2 is used for the separate aggregate transfer endpoint. See the Supplementary Material for architectures, teacher configurations, masks, normalization and spatial-condition construction.

### 3.2. From attribute error to instance evidence

Average attribute error and instance correspondence answer different questions. For reference *a*_*k*_(*I*), continuous readout error is

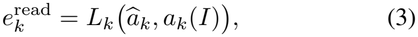

where spatial *L*_*k*_ is mean squared error and semantic *L*_*k*_ is one minus cosine similarity. Identification additionally specifies which alternative must be rejected. For sameconcept candidates *I*_*i*_, *I*_*j*_, a similarity score *S*_*k*_ gives the single-query margin

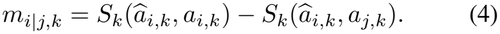

Its success credit is 1, 0 or 1*/*2 for positive, negative or tied margins. One query uses only the observation of its viewed image. Joint assignment instead uses both observations and tests the sign of *m*_*i*|*j,k*_ + *m*_*j*|*i,k*_. We report these tasks separately.

For squared distance, the dependence on the candidate difference is explicit. Writing *d* = *a*_*i*_ *−a*_*j*_, *µ* = (*a*_*i*_ +*a*_*j*_)*/*2, and 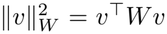 for fixed positive semidefinite *W*, the correct-candidate distance advantage is

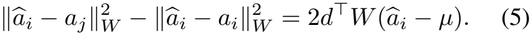

Equation 5 connects recovery error to the visual difference that determines an instance decision. Equal total errors can yield different decisions because their projections onto *d* differ. For each attribute, we characterize instance evidence by continuous readout error, matched-versus-exchanged correspondence and concept-level uncertainty. This profile is conditional on the participant, observation protocol, frozen representation, readout and scoring rule. The supported range comprises attributes and distinctions that meet the declared comparison and error criterion. The same coordinates enable downstream tracing and supply error targets for the risk model. Derivations of the candidate projection, joint-assignment cancellation, isotropic reduction and region-based decision conditions are given in the Supplementary Material.

### 3.3. Reliability as a testable evidence hypothesis

Once an instance decision is fixed, reliability asks which observations support it most strongly. We compare projected spatial-error models with ordinary margins, semantic agreement, full-feature confidence and wrong-correspondence controls. All scores rank the same spatial decisions.

For training-centered unit *R*_8_ candidates *v*_*A*_, *v*_*B*_ and prediction 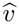, define

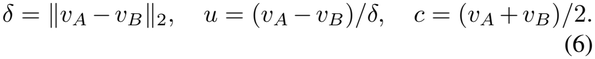

The sign of 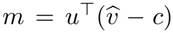 gives the common centeredcosine decision. Fittin g-concept residuals supply mean *µ* and regularized covariance Σ. The directional score is

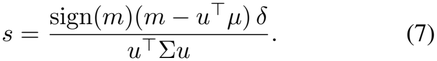

Shrinkage addresses covariance estimation stability [17]; its value for ranking remains an empirical question. This tests whether the projected error distribution helps rank correctness. Isotropic and diagonal variants test covariance structure; ordinary margins test simpler decision strength. Semantic agreement measures support from the separate pre-prior visual readout for the current spatial decision. Native semantic classification is reported separately. A learned full-feature head follows confidence-estimation practice [3]; cyclic and candidate-only scores test correspondence and candidate difficulty. Full definitions, covariance regularization, feature lists and degenerate cases are in the Supplementary Material.

## 4. Frozen Models and Evaluation Protocol

The historical panel uses project-preprocessed THINGS-EEG2 [9]. Sub-08 has 163 development concepts with ten images each, four-presentation EEG means, and 90 directed decisions per concept (14,670 queries). TRAIN and DEV images and concepts are disjoint, but if both historical fourslot selections came from the audited event archive, at least 6,462 of 6,520 nominal DEV 0–1000 ms windows intersect a TRAIN window in the same recording segment for every eligible slot choice. The exact historical selections and whitening membership are unknown. Candidate windows have zero overlap at 0–160 ms (Supplement Sec. 10).

Our primary recovery and reliability conclusions use the short-window evaluation. We hold out one physical session at a time and fit Ridge readouts on the other three, including TRAIN-only scaling. The paired recovery test uses the same event slots in [*−* 160, 0) and [0, 160) ms and averages two test presentations per image. The primary held-out reliability comparison uses sub-08’s post-onset test means and separate Ridge readouts for spatial geometry and CLIP semantics. The historical panel has different windows, observation averages and readouts. Its results cannot isolate the effect of session separation.

### Common decisions and matched retention

Selective classification evaluates error at retained coverage [8]. For the historical panel, five concept folds separate evaluation from error-statistic and confidence-head fitting. For the held-out panel, error models use concept-cross-fitted residuals from TRAIN. Every method retains half the queries within each concept, giving 7,335 retained decisions at the primary endpoint. Confidence ties are resolved independently of labels. This fixes both the decision rule and the concept-wise retention budget, so accuracy differences measure selection rather than a changed classifier. Complete risk–coverage curves are secondary.

### Paired uncertainty and main comparisons

Whole-concept bootstrap resampling preserves shared images, candidates and paired conditions. The four original historical directional-risk contrasts use joint multiplicity adjustment against isotropic error, semantic agreement, the full head and cyclic evidence. Other table intervals describe individual scores. Intervals condition on fitted models; the exposed development setting remains exploratory. Exact fold construction, draws, tie rules and numerical checks are provided in the Supplementary Material.

### Reproducibility

The supplement documents split construction, feature coordinates, scoring equations, and reproduction checks. The analysis uses existing public datasets; the supplement identifies the sources and the protocols used with each.

## 5. Results: Recovery, Realization and Reliability

### 5.1. Recoverability: instance detail survives in the readout

#### Short-window recovery with sessions held out

On 19,560 paired event slots across three participants, postonset matched *R*_8_ gains over cyclic exchange are +3.46, +3.74 and +3.26 percentage points for sub-08/01/02. Gains over coarse interpolation are +2.62, +3.20 and +2.66 points. All six post-onset and five of six post-minuspre intervals exclude zero, using 18 simultaneous concept intervals. All six pre-onset gain intervals include zero. The preceding stimulus and sequence effects can still contribute to short-window EEG (Fig. 3; Supplement Sec. 11).

**Figure 3.**
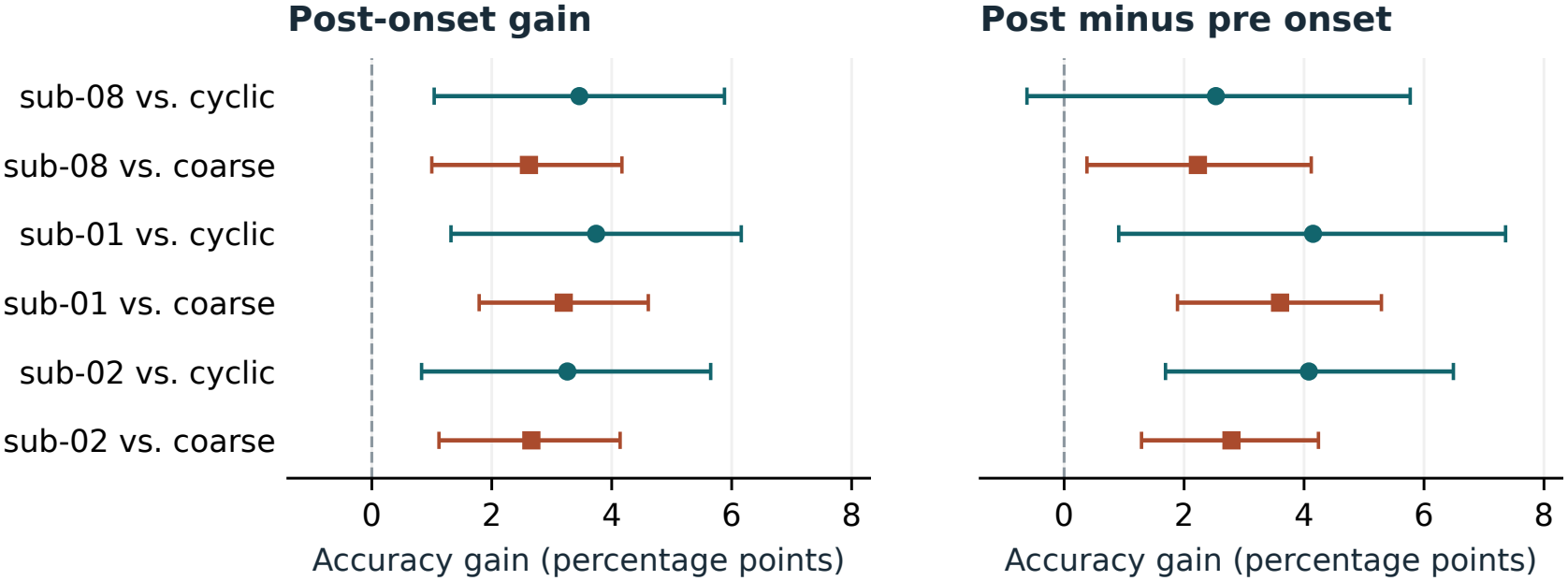
Fine spatial detail is recoverable in short windows with physical sessions held out. Left: matched *R*_8_ accuracy minus same-concept cyclic exchange or coarse interpolation after onset. Right: the change in that gain from [*−* 160, 0) to [0, 160),ms on the same events. There are 6,520 event slots per participant, two presentations per held-out image, and 163 resampled concepts per participant. Bars are 95% simultaneous intervals from the 18 original contrasts. The full pre-/post-onset estimates, fit scope and fold counts are in Supplement Sec. 11.

#### External recovery tests

Neither historically tested THINGS-EEG2 participant, sub-03 or sub-10, clears both controls. With the same 32 concepts and 64 single-presentation images per person, none of nine new NOD participants clears both controls for either tested readout. The four simultaneous group intervals all contain zero and the +3-point target. These results leave generalization across participants or recording protocols unresolved. Table 1 summarizes the scopes of these comparisons. Supplement Secs. 5–7 give the full scale profiles and external controls.

**Table 1.** Recovery evidence by dataset and protocol. Each row summarizes its own evaluation. The two shared-gallery comparisons use a matched gallery across datasets. The supplement also reports category, semantic instance, *D*_4_, and *R*_16_ evidence where measured.

| Dataset | Evaluation protocol | People | *R*<sub>8</sub> finding | Supplement |
|---|---|---|---|---|
| THINGS-EEG2 | Within person; physical session held out | 3 | Above cyclic and coarse after onset | Sec. 11 |
| THINGS-EEG2 | Cross-person, shared gallery; four presentations | 7 | Below coarse in point estimates | Sec. 5 |
| Alljoined-1.6M | Cross-person, shared gallery; four presentations | 7 | Below coarse in point estimates | Sec. 5 |
| NOD-EEG | Frozen THINGS readouts; single presentation | 9 | Group comparison intervals include zero | Secs. 6–7 |

#### Joint assignment can conceal single-query difficulty

The original sub-08 *R*_8_ predictions achieve 64.54% joint assignment but 52.73% single-query identification with squared distance (Table 2). A candidate-dependent preference common to two queries can cancel in joint assignment. The fixed-reference decomposition finds a strong energy-related component of this bias. Changing only the score raises single-query accuracy to 59.36% with all prediction arrays unchanged. The gain therefore reflects the scoring rule, not a newly trained encoder. The same fixed rule yields a 6.10-point single-query gain on the historical 100-concept panel (see the Supplementary Material).

**Table 2.**
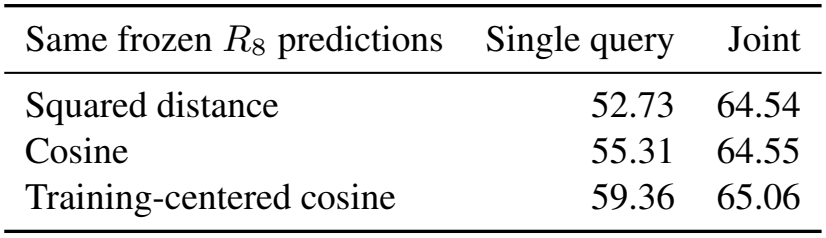
The scoring rule changes apparent capability. Accuracy (%) on sub-08, 163 development concepts, 1,630 images, 7,335 pairs and 14,670 query directions. Each EEG observation averages four presentations. Evaluation uses DA3 references; centering uses the original DAv2 training template.

| Same frozen *R*<sub>8</sub> predictions | Single query | Joint |
|---|---|---|
| Squared distance | 52.73 | 64.54 |
| Cosine | 55.31 | 64.55 |
| Training-centered cosine | 59.36 | 65.06 |

#### Added detail exceeds fixed interpolation

Historical DA3 *R*_8_ joint-assignment gains over exchange are 10.35,10.63,5.62 points for sub-08/01/02 on 100 exposed concepts. On the separate 163-concept DEV panel, single-query gains over fixed bilinear *D*_4_ interpolation are 5.60,3.14,1.47 points, with positive simultaneous intervals. These test the declared coarse explanation; *R*_16_ effects remain unresolved.

#### Frozen representations support additional visual recovery

On the exposed development panel, separately fitted probes recover *R*_8_ detail in three participants (Table 3, Fig. 4). MLP matched-minus-cyclic gains are 11.49 [8.73,14.22], 6.67 [4.14,9.26] and 3.71 [1.61,5.85] points for sub-08, sub-01 and sub-02; gains over fixed coarse interpolation are 7.79 [6.16,9.40], 3.59 [2.12,5.05] and 2.57 [1.13,4.02]. Linear-versus-MLP intervals include zero for all three. Comparing readout families tests probe dependence [11]. These comparisons support within-participant recovery. They do not test cross-person model transfer or isolate physiological differences. Training coverage and eligible-presentation averages differ, while every DEV observation averages four presentations.

**Table 3.** Within-participant recovery replicates. Training-centered cosine *R*_8_ single-query accuracy (%), averaged over two fitting seeds. Each participant has the same 163 exposed DEV concepts, 1,630 images and 14,670 query directions from four-presentation means. Coarse uses fixed bilinear interpolation of the predicted parent. Probes are fitted separately; training coverage differs.

| Participant | Probe | Matched | Cyclic | Coarse |
|---|---|---|---|---|
| sub-08 | Linear | 61.59 | 49.86 | 53.14 |
| sub-08 | MLP | 61.23 | 49.74 | 53.44 |
| sub-01 | Linear | 55.85 | 48.35 | 51.70 |
| sub-01 | MLP | 55.24 | 48.57 | 51.65 |
| sub-02 | Linear | 54.00 | 50.45 | 50.83 |
| sub-02 | MLP | 53.48 | 49.77 | 50.91 |

**Figure 4.**
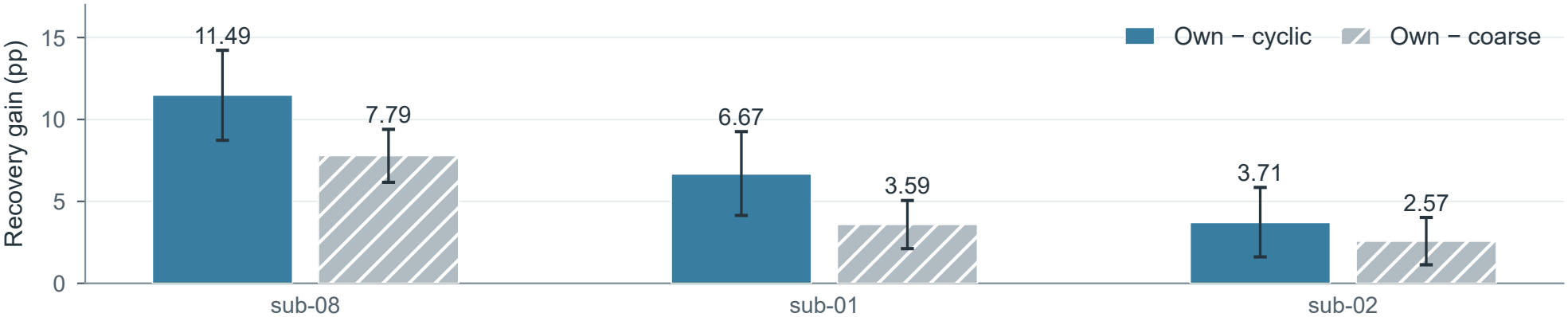
Recovered detail distinguishes same-concept instances. MLP *R*_8_ single-query gains over same-concept exchange and fixed coarse interpolation, with the original within-participant intervals. Each participant: 163 exposed concepts, 1,630 images, four-presentation means, and two probe seeds. Separately fitted probes have different training coverage. These original comparisons are distinct from the broader simultaneous scale-profile analysis.

#### Observation protocol

Trial averaging affects time-resolved decoding [10]. The primary results use four-presentation means. Mean-trained probes retain weaker correspondence on individual presentations; complete comparisons and protocol-shift analysis are in the Supplementary Material.

#### Semantic alignment

The historical pre-prior representation gives 75.84% DEV single-query identification versus 47.36% same-concept exchange on sub-08 four-presentation means. This motivates a full-representation quality baseline alongside spatial risks. The Supplementary Material reports complete scale- and seed-level recovery results, single-presentation comparisons, appearance controls and additional recording-condition analyses.

### 5.2. Realization: input evidence and output fidelity diverge

We exchange detail within concept while holding recipient semantics, coarse layout and generation noise fixed. For sub-08, the comparison uses 128 stimuli from 32 concepts, 192 within-concept pairs and three noise seeds. The generator consumes 16 *×* 16 conditions. At the 128 *×* 128 DAv2 scoring grid, the Own-minus-Exchange joint-assignment advantage falls from +25.52 to +0.69 percentage points, a change of *−*24.83 [*−*41.90, *−*6.71] points (12 simultaneous intervals; Fig. 5). Rescoring changes the measurement grid for fixed inputs and outputs. Only the historical 8 × 8 analysis covers three participants.

**Figure 5.**
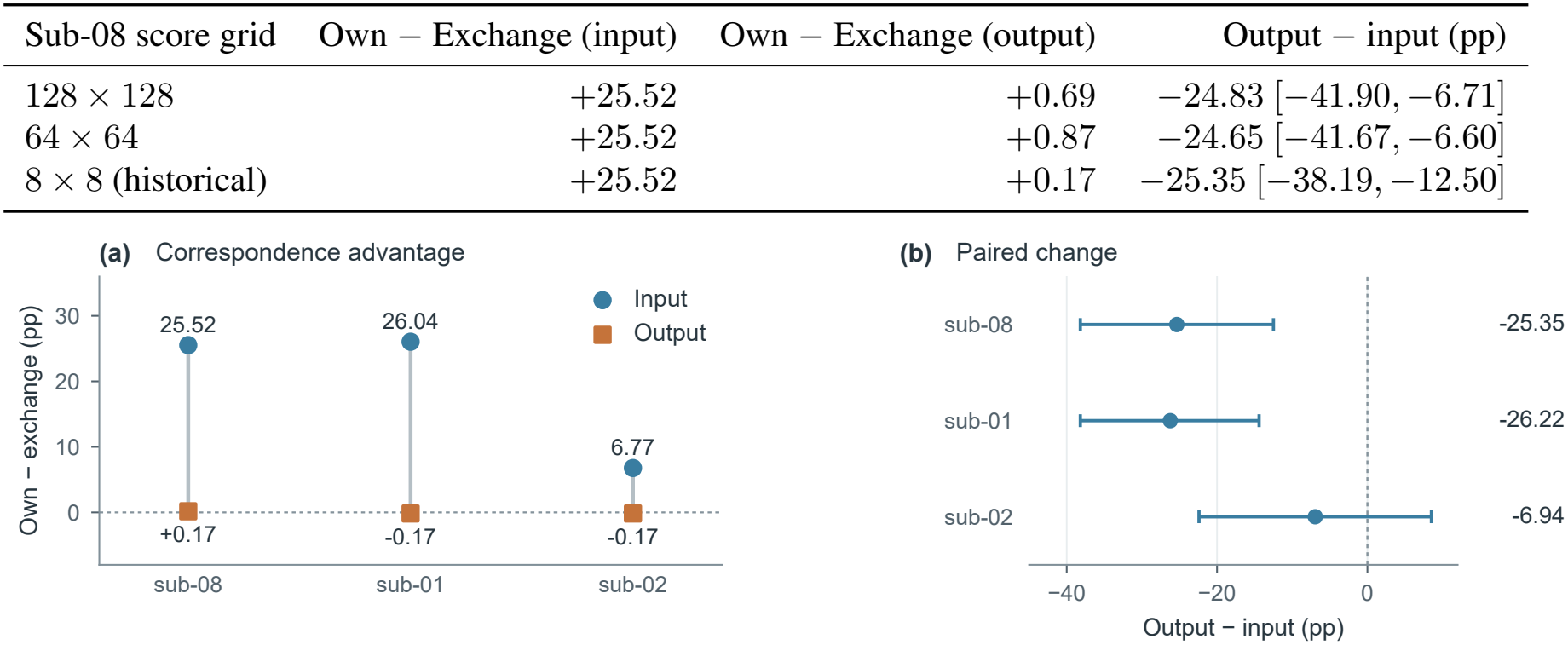
Joint depth assignment weakens after generation. Top: sub-08 rescoring of fixed outputs. The two finer-grid intervals use the 12-comparison adjustment. Bottom: historical 8 *×* 8 scores across three participants with marginal concept-bootstrap intervals. Controlled flips yield 22/24 versus 13/24 correct joint assignments at 128^2^ versus 8^2^, and both states are independently correct in 5/24 versus 0/24. At 128^2^, the equal-dose visual-input test yields *−* 1.39 [*−* 5.56, 0.00] pp of output advantage (eight adjusted intervals). The endpoint measures joint depth correspondence under the tested pipeline.

The finer-grid measurement orders known flips above chance jointly, but its independent state sensitivity remains limited. In the natural output panel, independent scoring gets both states right in 68/576 pairs in each arm at 128^2^, and the output-minus-input gain interval includes zero. In a separate 1,630-image single-query study, intervals for output advantage over the actual donor include zero for all three participants with either teacher (Supplement Sec. 1). The equal-dose visual-input test also gives no positive output gain, so the result does not isolate EEG as the cause. It remains compatible with loss in the generator interface and limits of output measurement. Natural-output visibility lacks a direct human test. Supplement Sec. 2 shows matched-scale controls, Sec. 3 gives the visual-input amplitude comparison, and Sec. 12 gives grid-specific calibration.

The separate sub-08 DA3 diagnostic uses 32 images from eight exposed concepts and one generation seed. Its historical cached scores give one positive adjusted input-minus-output interval in four contrasts (+18.75 [0.52, 38.54] points for the linear semantic readout versus cyclic detail). A fixed-teacher replay makes cached measurements repeatable. It does not validate this endpoint for human visibility. An eight-case latent trace finds six declines in location correspondence, with a mean change of *−* 0.097 [*−* 0.214, +0.032]. This exploratory result does not locate a stage of loss. The supplement reports the runtime injection checks, trace and full scoring controls (Sec. 12).

Separate EEG-only fidelity prediction and post-generation selection show no stable advantage for the tested structured risks or selection rule. Their complete comparisons and distinct label protocols are retained in the Supplementary Material.

### 5.3. Reliability: selection under session holdout

On the sub-08 short-window panel, the unfiltered spatial accuracy is 53.10% (Table 4). We retain 45/90 of the same decisions per concept for every score. At matched retention, signed semantic support improves over hash by +2.02 [1.04, 2.99] pp. Comparisons against plane margin (*−* 0.76 [*−* 1.99, 0.49] pp) and binary agreement (*−* 0.02 [*−* 0.57, 0.52] pp) remain unresolved (ten simultaneous concept intervals). Directional minus isotropic is *−* 0.17 [*−* 0.74, +0.40] pp, and directional minus diagonal is *−* 0.01 [*−* 0.52, +0.50] pp (eight-comparison adjustment). The learned head is unreported in this comparison because its fit emitted an optimizer warning.

**Table 4.** Selection at fixed coverage with sessions held out. Sub-08, 45/90 fixed spatial decisions retained per concept. Signed-support contrasts use the ten-comparison adjustment, and covariance contrasts use the eight-comparison adjustment. Supplement Secs. 9 and 11 give the additional ranks and treatment of the unresolved learned fit.

| Ranking score | Retained accuracy (%) | Paired contrast | Difference (pp) | Simultaneous 95% CI |
|---|---|---|---|---|
| Hash | 52.44 | Signed support − hash | +2.02 | [+1.04, +2.99] |
| Plane margin | 55.22 | Signed support − plane margin | −0.76 | [−1.99, +0.49] |
| Binary agreement | 54.48 | Signed support − binary agreement | −0.02 | [−0.57, +0.52] |
| Signed semantic support | 54.46 |  |  |  |
| Directional covariance | 55.08 |  |  |  |
| Isotropic error | 55.25 | Directional − isotropic | −0.17 | [−0.74, +0.40] |
| Diagonal error | 55.09 | Directional − diagonal | −0.01 | [−0.52, +0.50] |

#### Same-image donors test observation specificity

In a transfer diagnostic, THINGS Ridge shows Own minus same-image donor at 0.07 [0.48, +0.32] points while same-image minus other-image donors is +4.94 [3.47, 6.52]. Separate two-session holdouts give Own minus cross-session same-image donor of 0.06 [0.19, +0.06] and +0.03 [0.08, +0.15] points for sub-01/02 using a 2/4 quota. The comparisons are consistent with shared image difficulty and leave the added value of the current observation unresolved. The supplement reports a matched TrustScore-style gallery comparison [14] and the full donor controls (Sec. 14).

#### Historical development panel

At full coverage, every method uses the same base accuracy, 61.22%, with zero unresolved queries. Differences in Table 5 therefore concern correctness ranking alone. Native semantic classification scores 75.84% on all queries; semantic agreement instead selects among the fixed spatial decisions.

**Table 5.** Historical development panel at matched retention. Percentages at 45/90 retained directed queries per concept, 163 concepts and 7,335 retained queries. All ten scores retain the same fixed spatial base decisions. Table intervals describe each method, not the four family-adjusted differences.

| Confidence score | Accuracy | Risk [95% CI] |
|---|---|---|
| Spatial error models |  |  |
| Directional covariance | 66.41 | 33.59 [31.36,35.88] |
| Isotropic error | 66.94 | 33.06 [30.84,35.35] |
| Diagonal error | 66.95 | 33.05 [30.77,35.38] |
| Scalar spatial margins |  |  |
| Absolute plane margin | 66.79 | 33.21 [31.03,35.43] |
| Absolute cosine gap | 67.08 | 32.92 [30.69,35.22] |
| Semantic and full-feature controls |  |  |
| Semantic agreement | 83.65 | 16.35 [14.56,18.19] |
| Absolute semantic gap | 61.85 | 38.15 [36.33,39.97] |
| Full correctness head | 74.67 | 25.33 [23.30,27.38] |
| Correspondence and candidate controls |  |  |
| Cyclic evidence | 60.23 | 39.77 [37.96,41.62] |
| Candidate geometry only | 61.64 | 38.36 [36.52,40.25] |

The covariance score retains 66.41% correct queries, versus 66.94% isotropic, 67.08% ordinary cosine gap, 83.65% semantic agreement and 74.67% full-head confidence. Its risk differences are +0.53 [*−* 0.11,+1.17] points versus isotropic, +17.25 [15.02,19.39] versus semantic agreement and +8.26 [5.96,10.55] versus the full head. Positive values favor the comparator. Against cyclic evidence, risk is lower by 6.18 [3.60,8.70] points. This comparison changes image correspondence and image difficulty together. It therefore cannot establish an advantage over a different observation of the same image.

A separate nested concept-fold comparison controls semantic agreement, ordinary cosine gap and candidate distance with equal-capacity four-feature heads. The structured head retains 84.16% correct decisions; replacing its fourth feature with ordinary margin, isotropic score, cyclic score or candidate geometry gives 84.17–84.29%. All four paired intervals include zero. Agreement-based selection performs well, but these controls do not support an additional benefit specific to directional covariance. The Supplementary Material gives all nested-control contrasts and paired intervals.

The five principal curves in Fig. 6 retain the same decisions at every count. At 24.44% and 74.44%, directionalscore accuracies are 71.03% and 63.61%. These are secondary operating points, not certified probabilities or retrospectively optimized thresholds. No method or endpoint replaces the prespecified 50% covariance comparison after inspecting these results.

**Figure 6.**
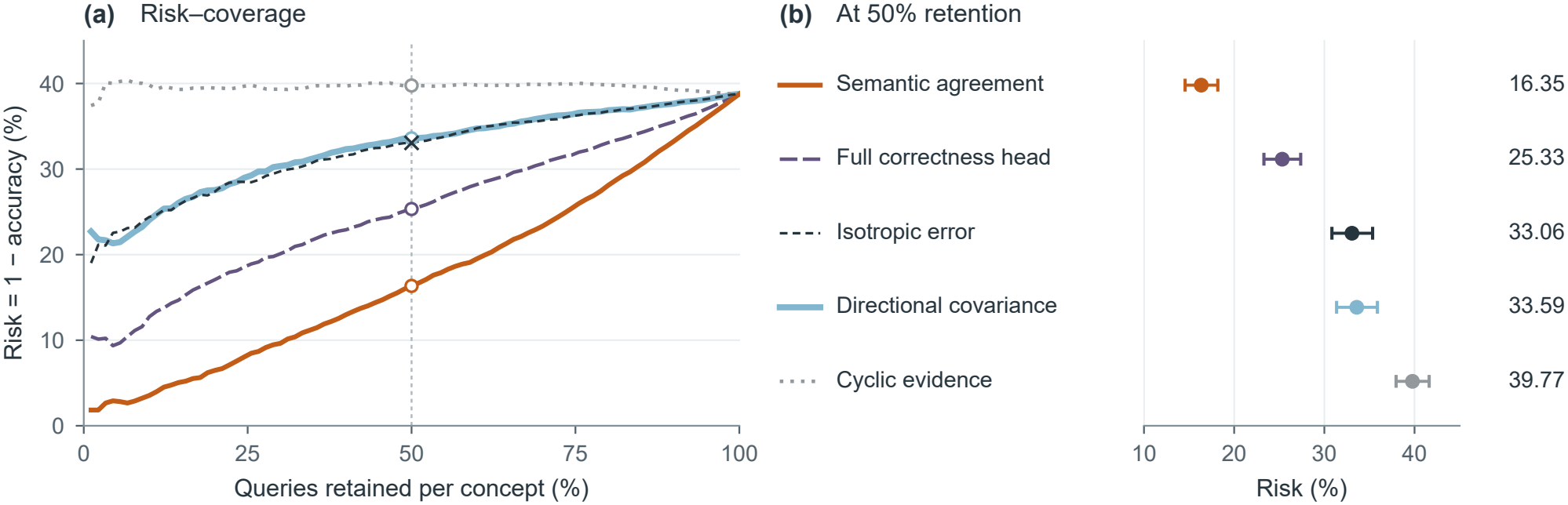
Historical development-panel risk–coverage curves. Five prespecified scores rank the same spatial decisions. (a) Recorded risk–coverage curves with equal retained counts per concept. (b) Risk at 50% retention (45/90 queries per concept), with saved marginal 95% intervals rather than paired risk-difference intervals. Colours and line styles identify the same scores in both panels. Semantic agreement ranks spatial decisions rather than performing native semantic classification.

### 5.4. Additional historical development panels

The same fixed-score, five-fold protocol evaluates sub-01/sub-02 with nine available scores after full-head fitting reaches its prespecified limit (Table 6). Directional-minus-isotropic risk intervals include zero for both participants, and directional risks exceed semantic-agreement risks. The cyclic contrast remains unresolved in sub-02. These conditional replications retain the exposed-data setting and participant-specific fits. Complete participant-specific curves, retention summaries and failed-fit records are included in the Supplementary Material.

**Table 6.**
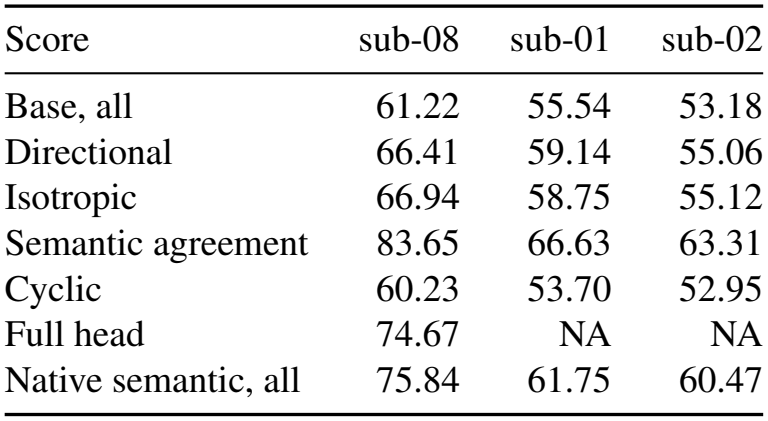
Secondary replications retain the strong-control limitation. Percent accuracy. Each participant has the same 163 concepts, 1,630 images and 14,670 four-presentation-mean queries. The middle scores retain 45/90 common spatial decisions per concept; the base and native semantic rows classify all queries. Probes and residual statistics are participant-specific. Full-head fits did not converge for sub-01/sub-02 and are reported as NA; detailed stopping records are supplementary. Only sub-08 supplies the full ten-method comparison.

## 6. Discussion and Limitations

Reconstruction quality combines three questions that our results examine separately: whether an attribute can be recovered, whether an instance decision is reliable, and whether the output preserves the distinction. Signed semantic support improves over hash in the session-heldout panel, with gains over simple spatial margins and binary agreement unresolved. The large historical ranking effect motivates these comparisons but cannot establish independent-session or observation-specific reliability. The comparison evaluates the fitted scores and does not isolate neural noise or establish a universal failure of covariance estimation. Perceptual realism and reference fidelity are distinct objectives [1]. Here, correspondence attenuation concerns the tested synthesis and evaluation chain; interfaces, semantics and output measurement remain possible causes.

### What a reconstruction evaluation must distinguish

The fixed-prediction comparison shows why one aggregate score cannot define the information recovered from EEG. A decision rule can emphasize shared candidate preferences or the direction that separates instances. Reporting the candidate task alongside attribute error makes the distinction explicit: a lower error and a more identifiable instance are different outcomes. This motivates reporting both individualquery decisions and joint assignments when evaluating a new decoder.

The generation comparison adds a second requirement. An output that changes after a condition exchange establishes sensitivity; faithful realization additionally requires that the change preserve the intended instance distinction. Comparing the same attribute before and after synthesis makes this requirement testable. The observed attenuation directs attention to the complete transfer path, including how spatial conditions are represented, injected and evaluated, rather than treating visible output change as sufficient evidence of control.

For reliability, the informative signal is support for a particular existing decision. Semantic agreement asks whether another visual readout supports that decision; an absolute semantic margin asks how strongly the semantic readout prefers either candidate. Their different roles explain why a semantic classification result cannot replace a commondecision ranking test. Matched retention then measures selection value while holding concept composition fixed. Together, these checks provide a practical way to assess a reconstruction system without collapsing recovery, responsiveness and correctness into one notion of quality.

The historical ranking panel uses development data with possible overlapping recording samples under the documented archive assumption. Concept cross-fitting cannot remove upstream exposure. Held-out results use refitted models with short windows and participant-specific data; they support conclusions at those settings. Bootstrap intervals condition on fitted models and recorded selections. External tests do not establish cross-person transfer or stable physiological limits. Monocular depth and semantic similarity measure declared feature spaces. Physical geometry and human fidelity require separate measurements.

Candidate references are legitimate side information for this task. True identity and current error remain offline labels; a score’s candidate context must not be described as gallery-free inference. Generation controls additionally depend on donor context, unlike the separate EEG-only fidelity heads. The relationship between recovery and useful output depends on both the generator and the candidate label. Outcomes under those conditions cannot certify the present risk ranking. The Supplementary Material provides the full public-benchmark comparison and reproduction records, alongside exploratory spatial-map and temporal controls.

## 7. Conclusion

Short-window session holdout supports within-participant spatial recovery for the tested distinctions. In the same framework, simple reliability controls narrow the interpretation of semantic ranking, and calibration bounds what downstream joint depth scores establish. Keeping these tests separate makes clear which neural evidence is recoverable and which claims about observation-specific reliability or visible fidelity remain open.

## Data Availability

The EEG analyses use three existing public datasets: THINGS-EEG2 (OSF: 3jk45), Alljoined-1.6M (dataset repository), and NOD-EEG (OpenNeuro: ds005811). Dataset-specific evaluation subsets and split rules are described in the Supplementary Material.

