## Supplement Material for "From Recoverability to Fidelity: Structured Visual Evidence Modeling for EEG Image Reconstruction"

Numbered citations refer to the main manuscript's References.

| Main result | What the supplement checks | Go to the evidence |
|---|---|---|
| Fig. 3, Table 1 | Sub-08 long-window overlap persists across four-slot choices in the archive. With sessions held out, five of six gains rose after onset at matched window lengths. | Overlap (Sec. 10), holdout (Sec. 11) Dataset summary (Sec. 5) |
| Figs. 1, 5 | Fig. 1 traces two same-category instance choices. On a separate panel, the joint depth advantage falls by about 25 pp after generation. | DA3 replay (Sec. 12) Fig. 5 scores (Sec. 12) |
| Table 4 | On held-out sessions, signed support beats hash. Gains over plane margin and binary agreement remain unresolved. The development-panel 83.65% is historical. | Held-out ranks (Sec. 9) TrustScore, donors (Sec. 14) |

#### 1. Do generated images retain instance detail?

Own uses the recipient's fine detail. Exchange takes it from another image of the same concept. Semantics, coarse layout and seed stay fixed.

| | | All nine candidates | | | Actual donor only | | |
|---|---|---|---|---|---|---|---|
| Participant | Reference | Input | Output | In. − out. | Input | Output | In. − out. |
| sub-08 | DAv2-Small | 10.19 | $-0.07$ $[-0.36, 0.21]$ | 10.27 $[7.62, 12.93]$ | 18.28 | $+0.43$ $[-0.49, +1.35]$ | $+17.85$ $[+12.26, +23.50]$ |
| sub-01 | DAv2-Small | 6.86 | $-0.14$ $[-0.49, 0.16]$ | 7.00 $[4.47, 9.56]$ | 11.84 | $+0.12$ $[-0.55, +0.86]$ | $+11.72$ $[+6.56, +16.69]$ |
| sub-02 | DAv2-Small | 2.64 | 0.20 $[-0.01, 0.41]$ | 2.44 $[0.58, 4.23]$ | 5.46 | 0.00 $[-0.61, +0.61]$ | $+5.46$ $[+1.35, +9.70]$ |
| sub-08 | DA3 | 7.59 | $-0.10$ $[-0.40, 0.21]$ | 7.68 $[5.00, 10.40]$ | 14.79 | $+0.06$ $[-0.80, +0.93]$ | $+14.72$ $[+9.45, +19.94]$ |
| sub-01 | DA3 | 5.53 | 0.16 $[-0.12, 0.43]$ | 5.37 $[3.13, 7.63]$ | 7.48 | $+0.49$ $[-0.37, +1.35]$ | $+6.99$ $[+2.32, +11.66]$ |
| sub-02 | DA3 | 1.08 | 0.03 $[-0.18, 0.25]$ | 1.05 $[-0.71, 2.83]$ | 3.25 | $-0.06$ $[-0.74, +0.61]$ | $+3.31$ $[-0.55, +7.19]$ |

**Full development panel.** Each person and condition has 1,630 images and 14,670 comparisons (1,630 actual donors). All donor output intervals include zero. Five win-rate and six margin gap intervals are positive (simultaneous 95% concept intervals, in pp). All-candidate output bounds are within $\pm 0.50$ pp [20, 28]. See Sec. 12 for the separate 128-stimulus Fig. 5 panel.

| Known occlusion order | Response direction | Image-state choice | Sensitivity |
|---|---|---|---|
| DAv2-Small (original score) | 11/24 | 24/48 | Unestablished |
| DA3 (original score) | 7/24 | 25/48 | Unestablished |
| Marked-strip color continuity | 24/24 | 48/48 | Synthetic crossing only |
| Two independent readers | 24/24 each | – | Visibility criterion met |

In 24 calibration-held-out concepts, global/local RGB ratios were .945/.883 (target) and .992/.979 (control). Material swap leaves continuity at 24/24 and reduces brightness to 0/24.

**Isolated flip.** Per-image amplitudes match in 24/24 pairs (fixed: 5/24). At $500^2$, target minus absolute distant gain is $+.0638$ $[-.0005, .2508]$ (DAv2-Small), $+.0129$ $[-.0007, .0420]$ (DA3), and $+.0102$ $[+.0006, .0318]$ (DA3 zoom $1.6\times$). Zoom–full and per-image–fixed intervals include zero. RGB arm means resolve both states in 15/24. Boundary scores give 22/24 at $512^2$ and 21/24 at $500^2$. At $512^2$, the marked flip ranks above each control in 24/24 (simultaneous lower bound 78.6%). Depth paired intervals include chance, with both states resolved in 0/24.

**Source photographs and natural outputs.** Readers passed 18/20 source-photo pairs. On six photos, both teachers beat same-surface controls in 2/6 at $16^2$ and 3/6 at $500^2$. The planned natural-output rating batch closed after qualitative category mismatch and produced no item-level judgments. Of 24 donor output intervals for $D_4/R_{16}$, only the sub-02 DA3 $D_4$ win rate excludes zero ($+0.67$ pp $[+0.06, +1.41]$). Its margin interval includes zero.

#### 2. Same-stimulus reconstructions and controls at matched scales

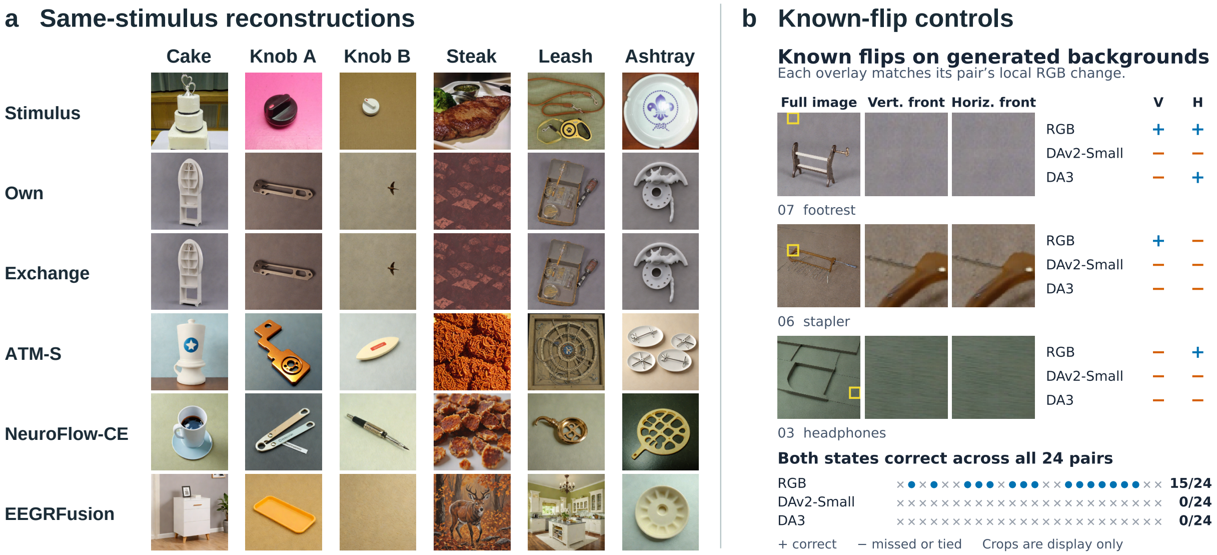


(a) Each THINGS-EEG2 stimulus has five outputs. EEGRFusion produced a nearly uniform image for knob B. (b) Three known-flip examples and scores for all 24 pairs. DEV48 has 24 concepts. CLIP and SwAV use the same 1,104 cross-class and 24 same-class image pairs.

| Output | Top-1 | Single | Both | CLIP cross | CLIP same | SwAV cross | SwAV same |
|---|---|---|---|---|---|---|---|
| Own | 4/48 | 29/48 | 2/48 | 74.73 | 14/24 | 87.77 | 19/24 |
| Exchange | 4/48 | 28/48 | 2/48 | 75.00 | 14/24 | 87.95 | 19/24 |
| ATM-S | 1/48 | 28/48 | 1/48 | 76.18 | 17/24 | 88.86 | 18/24 |
| NeuroFlow-CE | 1/48 | 26/48 | 1/48 | 72.37 | 16/24 | 75.18 | 13/24 |
| EEGRFusion | 2/48 | 26/48 | 1/48 | 74.73 | 14/24 | 89.40 | 18/24 |
| Ridge | 0/48 | 27/48 | 0/48 | 79.35 | 16/24 | 87.23 | 15/24 |
| Retrospective per-image selection | 6/48 | 39/48 | 4/48 | – | – | – | – |
| Original stimulus check | 44/48 | 48/48 | 44/48 | 100.00 | 24/24 | 100.00 | 24/24 |

Cross rates are percentages. CLIP scores the three single-output columns and the outcome-dependent selection. All CLIP cross intervals exceed chance. All CLIP same-class intervals include chance. For SwAV, 6/6 cross intervals exceed chance and 4/6 same-class intervals include it. 6/6 paired same-class differences include zero (20 simultaneous intervals). SwAV scores 24/24 after $32^2$ downsampling of originals.

**Official 200 (sub-08).** Compare matched and swapped CLIP labels in 19,900 pairs. Reference wins are 100% (194/200 top-1).

| | Geometry on | Geometry off | ATM-S | NeuroFlow-CE | EEGRFusion |
|---|---|---|---|---|---|
| Category top-1 | 8/200 | 6/200 | 16/200 | 13/200 | 8/200 |
| Assignment wins (%) | 80.62 | 77.64 | 84.03 | 79.08 | 80.10 |

All five rates exceed 50%. All four differences from geometry on include zero (simultaneous 95% intervals).

At $8\times8$, Spearman gains over unrotated other-image inputs are $+.874$, $+.894$ for known 0.5-pixel shifts and $+.016$ $[-.054, +.089]$ (24 pairs), $+.027$ $[-.057, +.114]$ (18 eligible) for generated pairs. Rotation gains are $-.061$, $-.073$. Median RGB change is 0.92 levels (8-bit); eligible cue boxes change 0.48 more than their surround.

**Signed texture response.** For 24 concepts, project luminance changes on two horizontal-shift templates. Known shifts are 0.5 pixels RMS. Coefficients of 1 indicate nominal amplitude (simultaneous 95% intervals).

| Shift direction | Known shift | Actual Own minus Exchange |
|---|---|---|
| Stimulus reference | $+1.125$ $[+1.089, +1.158]$ | $-0.002$ $[-0.045, +0.035]$ |
| Actual EEG input | $+1.158$ $[+1.131, +1.186]$ | $+0.013$ $[-0.010, +0.037]$ |

**Donor score at the observed $R_8$ angle.** After depth estimation, the matched input direction improves margins over actual and same-class other-image directions in 3/6 and 4/6 strata, and wins in 0/6 and 1/6 (simultaneous 95% intervals). The optimal direction can flip 89–132 of 1,630 pairs per stratum.

#### 3. Visual inputs at EEG-scale amplitudes

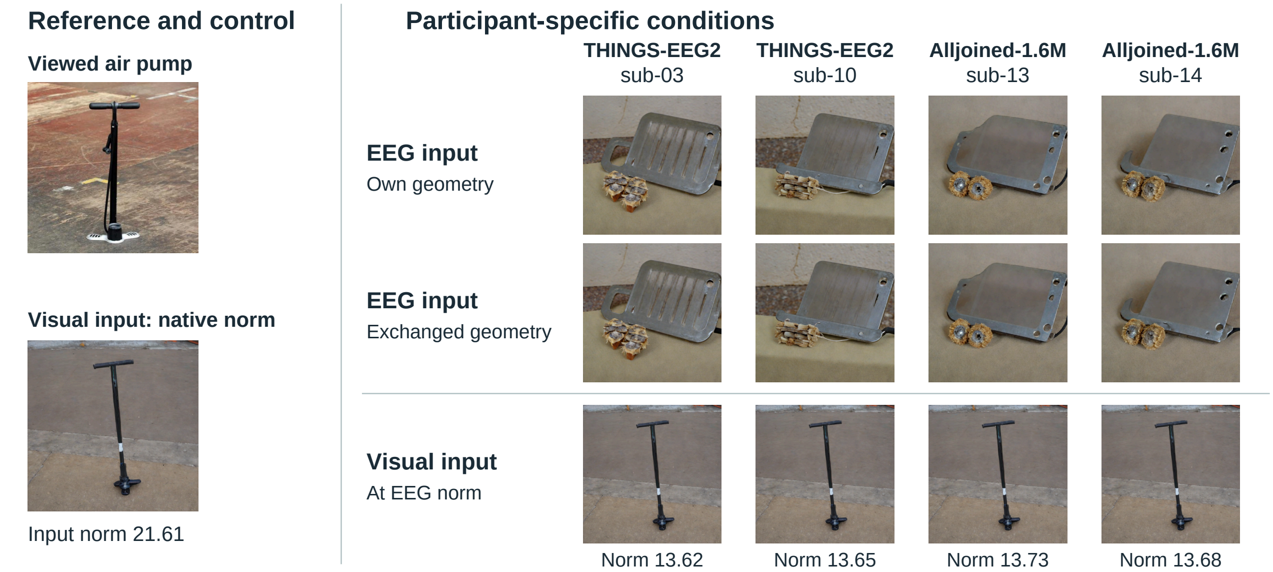


The gallery shows the second THINGS-EEG2 air pump. Within each column, compare Own with Exchange for EEG geometry. The bottom row uses the same visual input direction as the native-norm control, scaled to each participant's EEG norm. In a separate visual semantic-swap control for this stimulus, the output retained the category but favored the other air pump. The evaluation covers four held-out participants and both air pumps. EEG gives 0/8 pre-generation category hits. Changing vision norm alone (21.43 or 21.61 to 13.60–13.78) keeps 8/8 category (1,573 candidates) and instance (10) hits.

On eight other images, prior norm $1 \rightarrow 22.27$ raises the match margin $0.049 \rightarrow 0.060$ (2/4 concepts improve, preset 3/4), category $0/8 \rightarrow 1/8$, instance $5/8 \rightarrow 4/8$. Table changes are post-prior minus encoder on 1,630 sub-08 images, with unit vectors centered on 13,910 TRAIN references (eight simultaneous 95% intervals).

| Model | Instance pre / post (%) | Category change (pp) | Instance change (pp) |
|---|---|---|---|
| Project | 64.46 / 62.82 | $+0.80$ [$-1.29$, $+2.88$] | $-1.64$ [$-2.97$, $-0.40$] |
| ATM-S adaptation | 65.32 / 63.37 | $+2.33$ [$+0.03$, $+4.82$] | $-1.95$ [$-3.19$, $-0.74$] |

In EEGRFusion's native gallery, the prior loses 12 and gains 2 instance hits on the same 48 images ($-20.83$ pp, simultaneous 95% interval [-41.67, -2.08]). **EEGRFusion amplitude control.** Category, instance and joint hits across eight images are (6, 8, 6) for original vision, (3, 7, 3) for vision at EEG norm, and (0, 5, 0) for EEG. Both visual doses share direction and seeds. **Spatial adapter replay.** Across 32 images (eight concepts), gain is the same-image EEG/reference exchange cosine minus the mean for three same-concept alternatives. Intervals are simultaneous 95%.

| Stage | EEG/reference RMS | Same-image direction gain | Gain minus input gain |
|---|---|---|---|
| Input, $16\times16$ | – | $+0.240$ [$+0.012$, $+0.474$] | – |
| Adapter, $32\times32\times640$ | 0.996 [0.954, 1.044] | $+0.228$ [$+0.020$, $+0.431$] | $-0.012$ [$-0.060$, $+0.033$] |
| Adapter, $16\times16\times1280$ | 0.990 [0.949, 1.040] | $+0.218$ [$+0.004$, $+0.425$] | $-0.022$ [$-0.075$, $+0.025$] |

**Equal-amplitude outputs.** Over 32 images and three seeds, 12 depth contrasts span zero. Target and orthogonal shifts both reach $0.80$–$1.25\times$ the observed local RGB RMS in 84/96 reference and 77/96 EEG pairs. Those pairs cover 31/32 and 30/32 images, below the full-panel rule. Only 23/96 and 13/96 pairs match global RMS as well. On locally matched pairs, unit-dose target response minus orthogonal leakage is $+0.94$ [$+0.83$, $+1.03$] and $+1.00$ [$+0.95$, $+1.04$]. Archived target-axis estimates are $-0.0045$ [$-0.0182$, $+0.0018$] and $+0.0004$ [$-0.0034$, $+0.0038$] px (four simultaneous 95% intervals).

**Category-prompt intervention.** Six historical sub-08 inputs were selected because a frozen EEG readout ranked the target concept first. Each was rendered three times in CPU FP32 with the same EEG vector, geometry, seed and noise, changing only the empty, decoded or cyclic wrong-category prompt. Category / two-choice instance / joint hits were 1/3/1, 3/5/3 and 1/4/1 out of six per arm. The decoded prompt adds two category hits with no losses against either comparator. The descriptive paired interval for the gain is $[0, 66.7]$ pp. This selected panel shows a text-conditioning response. Full-gallery transfer and natural geometric validity are unresolved.

### 4. Does frozen decoding transfer across THINGS-EEG2 participants?

Each of ten THINGS-EEG2 participants contributes 1,630 images across 163 development concepts [9]. Four EEG presentations per image span two physical sessions. Six original training participants supply the sources. Evaluating one of those six leaves that person out. One-, two-, and multi-source Ridge fits share the images and training observation count within each split (6,240–7,050).

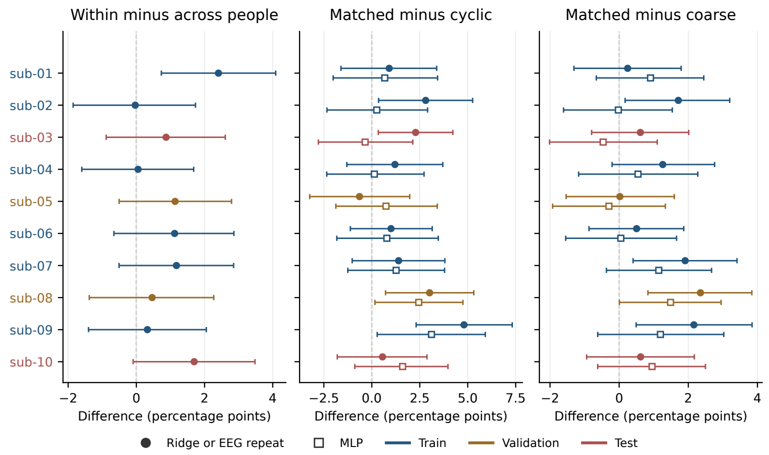


Multi-person Ridge and two-seed MLP use matched inputs and observations. Figure and per-participant table intervals are simultaneous over the planned contrasts within each participant and model. The source-count table equally weights participants, with descriptive 95% intervals that resample participants. Training cohorts overlap.

| Sources | Matched minus cyclic | Matched minus coarse |
|---|---|---|
| 1 | +0.62 [−0.21, +1.44] | +0.67 [+0.21, +1.12] |
| 2 | +1.18 [+0.63, +1.67] | +1.25 [+0.77, +1.65] |
| 5 or 6 | +1.73 [+0.75, +2.81] | +1.14 [+0.58, +1.71] |

Differences are in pp. One- and two-person fits use one fixed source composition per split.

#### 4.1. Per-participant fine-detail results with DAv2 references

| Person | Historical role | Sources | Ridge–cyclic | Ridge–coarse | MLP–cyclic | MLP–coarse |
|---|---|---|---|---|---|---|
| sub-01 | train | 5 | +0.91 [−1.61, +3.38] | +0.25 [−1.31, +1.79] | +0.67 [−2.01, +3.43] | +0.91 [−0.66, +2.45] |
| sub-02 | train | 5 | +2.81 [+0.35, +5.26] | +1.71 [+0.17, +3.20] | +0.26 [−2.34, +2.91] | −0.02 [−1.61, +1.54] |
| sub-03 | test | 6 | +2.28 [+0.34, +4.23] | +0.61 [−0.80, +2.01] | −0.35 [−2.78, +2.14] | −0.46 [−2.01, +1.10] |
| sub-04 | train | 5 | +1.21 [−1.30, +3.70] | +1.26 [−0.20, +2.76] | +0.13 [−2.35, +2.72] | +0.55 [−1.17, +2.27] |
| sub-05 | validation | 6 | −0.64 [−3.24, +1.98] | +0.02 [−1.53, +1.60] | +0.74 [−1.87, +3.41] | −0.30 [−1.93, +1.34] |
| sub-06 | train | 5 | +1.00 [−1.12, +3.15] | +0.50 [−0.87, +1.87] | +0.78 [−1.82, +3.47] | +0.05 [−1.55, +1.66] |
| sub-07 | train | 5 | +1.40 [−1.02, +3.81] | +1.91 [+0.40, +3.41] | +1.27 [−1.25, +3.79] | +1.15 [−0.37, +2.67] |
| sub-08 | validation | 6 | +3.01 [+0.72, +5.32] | +2.35 [+0.83, +3.84] | +2.45 [+0.16, +4.75] | +1.49 [+0.01, +2.95] |
| sub-09 | train | 5 | +4.81 [+2.31, +7.32] | +2.16 [+0.49, +3.84] | +3.11 [+0.28, +5.92] | +1.19 [−0.62, +3.03] |
| sub-10 | test | 6 | +0.56 [−1.80, +2.88] | +0.62 [−0.94, +2.17] | +1.61 [−0.89, +3.97] | +0.95 [−0.62, +2.49] |

Neither historically tested participant (sub-03, sub-10) clears both controls. The test participants and concepts had prior exposure in development.

**Within-person and cross-person Ridge.** For sub-01 and sub-02, both fits share 10,014 training images and score each participant on the same 7,335 pairs. Entries are joint minus single-query scores (pp) with three simultaneous 95% intervals conditional on these two participants.

| Endpoint | Within-person | Cross-person | Within minus cross |
|---|---|---|---|
| Matched score | +1.88 | +1.02 | +0.86 [−0.48, +2.30] |
| Gain over cyclic mismatch | +1.87 | +1.54 | +0.33 [−1.57, +2.31] |
| Gain over coarse interpolation | −3.51 | −0.14 | −3.36 [−5.77, −1.08] |

#### 5. How well do the readouts transfer between datasets?

**Recoverability by dataset and test protocol.**

| Test dataset and protocol | People (repeats) | Category | Semantic instance | $D_4$ | $R_8$ | $R_{16}$ | Details |
|---|---|---|---|---|---|---|---|
| THINGS-EEG2, within-person holdout by session | 3 (2 per session) | – | – | – | Exceeds both | – | Sec. 11 |
| THINGS-EEG2, cross-person, shared gallery | 7 (4) | – | 55.40/56.08% | Diagnostic | Below coarse | – | Sec. 5 |
| Alljoined-1.6M, cross-person, shared gallery | 7 (4) | – | 51.85/51.51% | Diagnostic | Below coarse | – | Sec. 5 |
| NOD, frozen THINGS readouts | 9 (1) | Unresolved | Unresolved | – | Unresolved | – | Secs. 6–7 |

Exceeds both means positive adjusted gains over cyclic and coarse. Below coarse refers to point estimates. Unresolved intervals span zero. $D_4$ tests repeats or sources. Dashes mark untested endpoints. Sec. 10 lists screened datasets.

**Matched two-dataset comparison.** Both datasets use 1,573 candidates and 738 queries from 159 concepts. Each has two source and eight held-out participants. Excluding one pilot per dataset leaves seven held-out participants per row. Semantic scores (Ridge/MLP above) use four-presentation means and the fixed training-image center. The other rows use separate protocols.

| Readout and endpoint | THINGS-EEG2 | Alljoined-1.6M | Difference (pp), 95% CI |
|---|---|---|---|
| Ridge semantic wins | 55.40% | 51.85% | $+3.55$ $[0.28, 6.50]$ |
| MLP semantic wins | 56.08% | 51.51% | $+4.57$ $[2.01, 7.15]$ |
| Ridge $R_8$ minus cyclic | $+0.59$ pp | $+0.85$ pp | $-0.26$ $[-4.23, 3.65]$ |
| MLP $R_8$ minus cyclic | $+0.48$ pp | $+0.48$ pp | $-0.01$ $[-3.88, 3.99]$ |
| Ridge $R_8$ minus coarse | $-2.27$ pp | $-1.45$ pp | $-0.82$ $[-3.82, 2.21]$ |
| MLP $R_8$ minus coarse | $-1.67$ pp | $-0.80$ pp | $-0.87$ $[-3.82, 2.01]$ |

Differences are THINGS-EEG2 minus Alljoined-1.6M. Intervals are simultaneous across the six comparisons. The semantic intervals are positive, and all four spatial intervals contain zero.

Domestic minus transferred semantic wins are $+5.27$ and $+6.04$ pp in THINGS-EEG2 and $-0.85$ and $-1.53$ pp in Alljoined-1.6M (Ridge and MLP, same participants).

**Semantic and spatial gaps under fixed weights.** The first four rows report THINGS-EEG2 minus Alljoined-1.6M (pp), with simultaneous 95% intervals across eight effects.

| Evaluation contrast | Ridge | Structured MLP |
|---|---|---|
| Same THINGS weights, four presentations | $+2.70$ $[-0.88, 6.32]$ | $+3.04$ $[-0.20, 6.55]$ |
| Same Alljoined weights, four presentations | $-1.72$ $[-4.49, 0.97]$ | $-1.46$ $[-4.43, 1.35]$ |
| Four minus single presentation, domestic weights | $+0.37$ $[-2.58, 3.34]$ | $+0.67$ $[-2.49, 3.87]$ |
| Matched minus cyclic, domestic weights, four presentations | $+3.05$ $[-0.70, 6.90]$ | $+4.87$ $[1.26, 8.72]$ |
| Semantic gap $-$ $R_8$ gap after cyclic subtraction (95% simultaneous intervals) | | |
| Domestic weights | $+3.31$ $[-2.24, 8.96]$ | $+4.88$ $[-0.25, 10.17]$ |
| Same THINGS weights | $+2.27$ $[-3.49, 8.12]$ | $+3.54$ $[-1.82, 9.13]$ |
| Same Alljoined weights | $-1.62$ $[-6.65, 3.45]$ | $-1.31$ $[-6.54, 3.95]$ |
| Paired source contrasts (simultaneous 95% intervals; 4 vs. 1 uses matched minus cyclic) | | |
| Matched: source interaction | $+4.42$ $[-0.11, +8.85]$ | $+4.51$ $[-0.09, +9.33]$ |
| Matched: equal-source mean gap | $+0.49$ $[-1.77, +2.81]$ | $+0.79$ $[-1.22, +3.05]$ |
| Matched minus cyclic: source interaction | $+4.63$ $[-1.43, +10.25]$ | $+5.43$ $[-1.09, +11.80]$ |
| Matched minus cyclic: equal-source mean gap | $+0.42$ $[-2.62, +3.59]$ | $+0.91$ $[-1.79, +3.59]$ |
| 4 vs. 1, equal-source gain gap | $+0.40$ $[-2.38, +3.10]$ | $+0.48$ $[-2.57, +3.58]$ |
| 4 vs. 1, change in source interaction | $+1.97$ $[-3.00, +6.79]$ | $+3.03$ $[-2.62, +8.77]$ |

All six semantic minus $R_8$ gap intervals include zero. The source interaction is the THINGS-weight minus Alljoined-weight gap. Weighting sources equally, the nearest and farthest $R_8$ candidates (2,920 directions each) give 24 simultaneous intervals spanning zero for match-minus-control gains and far-minus-near changes.

**Spatial gains over cyclic matching.** The plot compares four versus one presentation under domestic weights and domestic versus transferred weights for single presentations. All 16 simultaneous intervals include zero.

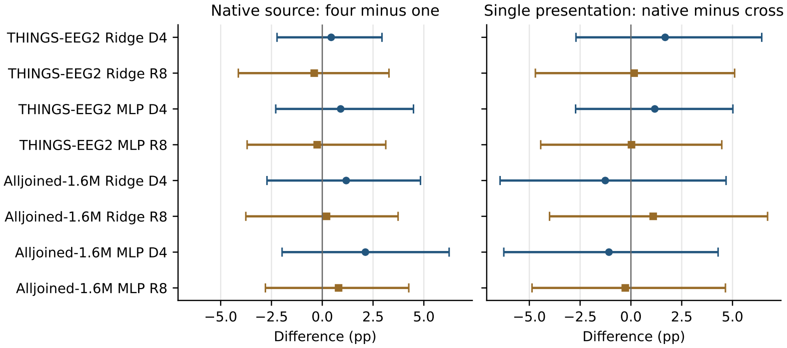

#### 6. Spatial decoding across nineteen NOD participants

THINGS Ridge and the two-seed MLP use 17 channels and four 40 ms bins from onset. We score matched $R_8$, same-concept EEG swap and $D_4$ interpolation against DAv2 references. The nine new participants each contribute 64 single-presentation images from the same 32 previously evaluated concepts. All 576 new images passed duplicate screening against the earlier panels.

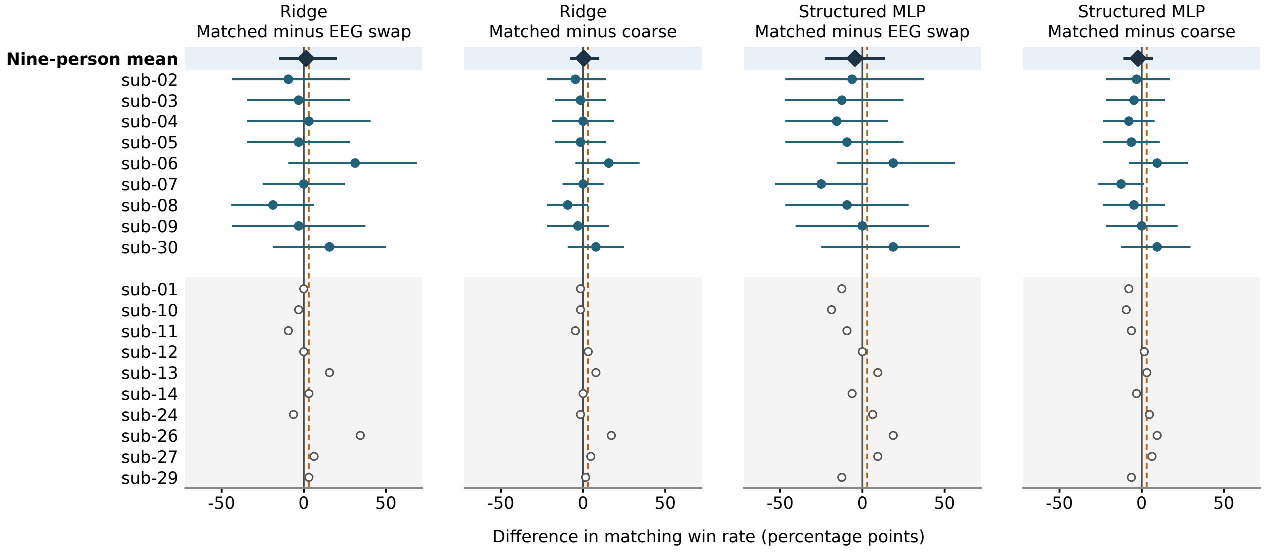


Filled circles show new participants and open circles show the historical ten as descriptive points. The diamond and table show the equal-participant mean for the new nine. Bars and brackets give simultaneous 95% intervals (36 individual contrasts and four group contrasts). The dashed line marks +3 pp.

| New nine participants | Matched minus swap (pp) | Matched minus coarse (pp) |
|---|---|---|
| Ridge | $+1.39$ $[-14.93, +20.14]$ | $+0.35$ $[-7.81, +9.72]$ |
| Structured MLP | $-4.51$ $[-22.57, +13.89]$ | $-2.26$ $[-11.11, +6.94]$ |

4/4 group intervals include zero and 4/4 include +3 pp. No new participant exceeded both controls for the same model by the simultaneous intervals.

**Matched training size and time controls.** We reuse all 384 NOD trials and the THINGS training center. Each reduced Ridge fits 696 trials from two people and 116 concepts (4 and 2 images per concept). THINGS and Alljoined share the same training images. NOD training holds out the evaluation participants and concepts. Full external fits use 14,012 trials each.

| Diagnostic contrast | Readout | Gain or change over mismatch | Gain or change over coarse |
|---|---|---|---|
| Alljoined − THINGS | Ridge | $-23.44$ $[-53.12, +6.25]$ | $-12.24$ $[-26.82, +2.60]$ |
| | Structured MLP | $-18.23$ $[-42.19, +5.73]$ | $-9.38$ $[-22.14, +2.35]$ |
| NOD gain | Ridge | $+0.00$ $[-13.54, +12.50]$ | $-0.52$ $[-7.81, +6.77]$ |
| NOD − THINGS | Ridge | $-9.38$ $[-36.46, +18.75]$ | $-5.47$ $[-19.01, +8.59]$ |
| THINGS 696 − full | Ridge | $-9.38$ $[-43.23, +20.83]$ | $-4.69$ $[-21.35, +10.94]$ |
| Alljoined 696 − full | Ridge | $+19.27$ $[-14.58, +55.21]$ | $+9.64$ $[-9.11, +28.91]$ |
| NOD − THINGS 696 | Ridge | $+0.00$ $[-25.26, +26.04]$ | $-0.78$ $[-15.24, +13.54]$ |
| NOD − Alljoined 696 | Ridge | $-5.21$ $[-34.38, +23.96]$ | $-2.86$ $[-19.40, +15.36]$ |
| Late − prestimulus | Ridge | $+0.52$ $[-23.96, 28.65]$ | $+0.26$ $[-11.98, 14.32]$ |
| | Structured MLP | $-8.33$ $[-30.73, 14.58]$ | $-5.21$ $[-16.67, 6.51]$ |

Table intervals include zero (four per family, eight for size matching). Late and pre are 80–160 and $[-100, -20)$ ms. In the new nine, shifting the 160 ms window by 40, 80 or 160 ms gives no established improvement over the original window (28 simultaneous 95% intervals).

#### 7. NOD controls for semantics, EEG signal and scoring

The nine new participants each use the same 64 images as in Sec. 6. The category score takes the top image in each person's gallery. Its control swaps EEG across concepts. The instance score compares the two images per concept against the within-concept swap. The last column subtracts the $R_8$ swap gain on the same images. Brackets give simultaneous 95% intervals across six contrasts.

| Readout | Category gain (pp) | Instance gain (pp) | Instance minus $R_8$ gain (pp) |
|---|---|---|---|
| Ridge | $-0.69$ $[-5.38, +3.12]$ | $+4.17$ $[-11.46, +19.44]$ | $+2.78$ $[-22.92, +25.35]$ |
| Structured MLP | $-0.35$ $[-3.99, +3.12]$ | $+4.86$ $[-11.11, +20.83]$ | $+9.38$ $[-15.97, +35.30]$ |

Both instance-minus-geometry intervals include zero. The initial two-person, 48-concept panel had four positive instance point estimates, with all eight semantic intervals including zero.

**Direct EEG signal control.** Each query has a same-concept reference from another run and a different-concept foil from that reference run. Windows $[-100, -20)$ and $[80, 160)$ ms share triples (29 eligible concepts in all nine new participants).

| Panel | Images | Concepts | Triples | Pre (%) | Late (%) | Late − 50% (pp) | Late − pre (pp) |
|---|---|---|---|---|---|---|---|
| sub-01, 100-concept panel | 400 | 100 | 14,328 | 48.61 | 50.12 | $+0.12$ $[-2.72, +3.09]$ | $+1.51$ $[-2.93, +5.87]$ |
| sub-10, 100-concept panel | 200 | 100 | 2,370 | 44.69 | 51.09 | $+1.09$ $[-5.93, +8.27]$ | $+6.40$ $[-3.16, +16.01]$ |
| Nine new participants | 576 | 29 | 1,065 | 47.32 | 51.69 | $+1.69$ $[-5.72, +8.96]$ | $+4.37$ $[-5.95, +15.62]$ |

Both nine-person concept intervals include zero. Across runs and concepts, EEG energy rises above pre-onset at 80–160 ms by $+1.36$ $[+0.33, +3.29]$ and $+1.09$ $[+0.44, +1.80]$ $\mu V^2$ in the six and nine participants, respectively, and rises at 160–240 ms (12 simultaneous comparisons). Within concepts, the product of cross-person EEG and DAv2 difference cosines gives eight $D_4/R_8$ intervals including zero (post and post minus pre, two cohorts, 960 images).

**Direction-injection control at fixed prediction norm.** For four earlier participants, mixing centered $R_8$ with matched DAv2 directions at weight 0.20 gave positive gains over both controls in seven of eight participant–model combinations. Full injection gave 100% matched wins. In the six-person extension (384 images), equal rotations toward the match or foil fixed the original prediction norm and pair. Zero rotation reproduced the archived scores.

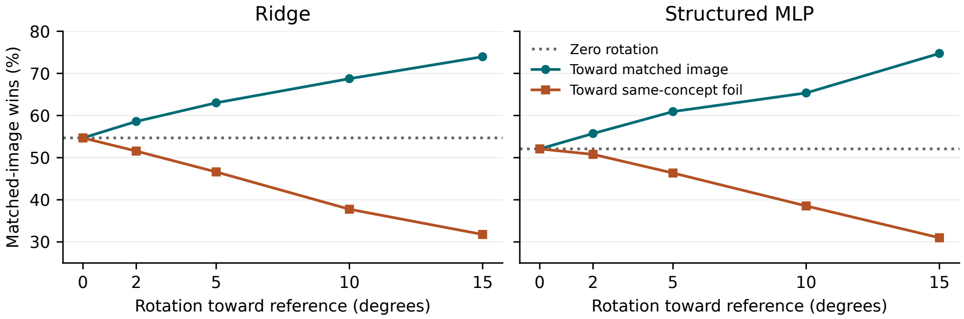


At 5°, matched-direction rotation exceeded foil-direction rotation by 16.41 pp [8.59, 25.78] for Ridge and 14.58 pp [7.29, 23.18] for MLP. Gains over zero rotation were 8.33 pp [2.60, 15.36] and 8.85 pp [2.86, 16.41]. Intervals are simultaneous across eight comparisons at 5° and 15°. These gains measure response to an injected reference direction.

**Single-presentation reliability.** In nine participants (32 previously evaluated concepts and 576 queries per readout), the Sec. 9 selectors keep one of two queries per person and concept. The first two contrasts compare signed support with agreement-then-spatial and absolute spatial margin at this fixed 50% coverage. The last contrast uses frozen THINGS training thresholds for signed support.

| Readout | vs. agreement (pp) | vs. spatial (pp) | Frozen kept | Frozen vs. full (pp) |
|---|---|---|---|---|
| Ridge | $+5.56$ $[-4.86, +16.32]$ | $+9.03$ $[-4.51, +22.92]$ | 277/576 | $+2.74$ $[-4.62, +10.50]$ |
| MLP | $+1.74$ $[-9.72, +13.19]$ | $+3.47$ $[-10.42, +16.67]$ | 245/576 | $+2.46$ $[-6.56, +11.24]$ |

All simultaneous gain intervals include zero, as in the initial two-person panel (48 concepts).

#### 8. Retrieval and spatial baselines on the same gallery

Tables share 1,630 sub-08 development images. Category top-1 uses the highest score per concept. Each query has nine same-concept comparisons, with half credit for ties. ATM-S adapts the author CLIP branch [18].

| Archived readout / reference | Category top-1 | Mean pairwise win rate |
|---|---|---|
| Project EEG-to-CLIP | 14.48% | 75.84% |
| Project after prior | 9.14% | 68.51% |
| ATM-S encoder (adaptation) | 15.21% | 76.16% |
| ATM-S after prior | 9.20% | 67.98% |
| NICE architecture, local fit | 10.37% | 71.36% |
| UBP encoder, local fit | 9.51% | 71.19% |
| Ridge-to-CLIP | 4.79% | 56.81% |
| Geometry reference from DA3 | | |
| Linear R8 | 1.50% | 61.59% |
| MLP R8 | 1.44% | 61.23% |
| Coarse interpolation | 0.52% | 53.44% |
| Cyclic R8 | 1.44% | 49.74% |
| Constant reference (ties) | 0.61% | 50.00% |

**Visual layer alignment on the same gallery.** Following the intermediate-target comparison in Shallow Alignment [6], we train two NICE readouts [24] for three epochs on 13,910 images. Both use L2-normalized means of the 256 patch tokens from the same CLIP ViT-H/14. Only the target block changes.

| Target block | Category top-1 | Within-concept wins | Within-concept top-1 | Full-gallery top-1 |
|---|---|---|---|---|
| 16 (middle) | 14.29 | 78.54 | 36.44 | 4.97 |
| 32 (final) | 7.30 | 69.37 | 24.97 | 2.09 |
| Difference (pp) | +6.99 [+3.62, +10.31] | +9.18 [+7.67, +10.76] | +11.47 [+8.71, +14.36] | +2.88 [+1.35, +4.42] |

Rates are in percent. Brackets give simultaneous 95% concept-bootstrap intervals. Cyclic within-concept wins are 47.21% and 47.70%. Projected-CLIP NICE has 48.08% cyclic wins.

**Geometry from frozen representations.** Identical Ridge probes fit DAv2 geometry on 13,909 training images. Each condition has 14,670 single-query comparisons or 7,335 joint assignments. Brackets give simultaneous 95% intervals for the four layer contrasts and the separate eight-contrast UBP family.

| Ridge input | Two-way (%) | Joint (%) | Ten-way (%) | Two-way − cyclic (pp) | Two-way − coarse (pp) |
|---|---|---|---|---|---|
| EEG (projected CLIP NICE) | 59.58 | 65.54 | 16.75 | +10.94 | +5.96 |
| EEG (UBP, fixed blur) | 59.62 | – | 16.50 | +10.20 [+7.10, +13.21] | +5.36 [+3.58, +7.19] |
| EEG (UBP, adaptive blur) | 59.72 | – | 16.44 | +10.32 [+7.21, +13.29] | +5.43 [+3.67, +7.27] |
| Image (projected CLIP) | 63.71 | 72.91 | 19.51 | +15.28 | +6.44 |
| EEG (block 16) | 60.54 | 66.09 | 17.79 | +11.42 | +6.45 |
| EEG (block 32) | 59.50 | 64.95 | 16.44 | +10.50 | +6.06 |
| Image (block 16) | 81.72 | 92.61 | 42.88 | +34.81 | +11.40 |
| Image (block 32) | 66.61 | 78.88 | 20.92 | +17.42 | +7.01 |
| Middle minus final (EEG) | | | | +0.91 [−1.32, +3.15] | +0.39 [−0.95, +1.75] |
| Middle minus final (image) | | | | +17.39 [+14.81, +19.96] | +4.39 [+2.59, +6.17] |

Joint scoring raises NICE matched scores by 5.45–5.95 pp and reduces coarse-adjusted gains by 2.95–3.08 pp. All six NICE gallery-change and 12 held-out intervals contain zero on the shared 1,500 pairs. For exact top-1, all five EEG readouts exceed both controls with 2, 4 and 6 candidates. With ten, all five exceed cyclic mismatch and two exceed coarse interpolation (40 simultaneous 95% intervals). Smaller sets average all subsets that include the target.

**UBP retrieval.** Author UBP [25] replays 114/200 top-1 and 169/200 top-5 (mAP 0.691769, RN50, 80 repeats). Its training pool includes DEV. Local UBP uses 17 electrodes and NICE 63. With 50 epochs and a shared medium-blur gallery, fixed and adaptive retrieval reach 76.71% and 76.65% within-concept wins [27], a difference of −0.06 [−0.22, +0.10] pp. Gain differences for adaptive versus fixed UBP and fixed UBP versus NICE include zero for both controls.

#### 9. Which rankings retain more correct decisions?

**Whole-session holdout (sub-08).** Keep 45/90 fixed decisions per concept. After onset, all decisions are 53.10% correct. Hash, plane margin, binary agreement and signed support retain 52.44/55.22/54.48/54.46% correct. Signed gains over the first three are $+2.02$ $[1.04, 2.99]$, $-0.76$ $[-1.99, 0.49]$ and $-0.02$ $[-0.57, 0.52]$ pp. Signed accuracy rises from 50.05% before onset by $+4.40$ $[2.46, 6.41]$ pp (ten simultaneous intervals). Fit geometry/CLIP Ridge on TRAIN and average two test presentations. See Sec. 11 for more rankings.

**Historical development panel (sub-08).** With different windows, readouts and repeats, signed support retains 83.65% correct at 45/90, versus 81.02% for binary agreement with hash ties and 81.55% for its fixed-marginal null. The null shuffles semantic correctness within concept, preserves both correct counts and the quota, and averages agreement-first selection over random ties. The gains are +2.63 [2.03, 3.29] and +2.11 [1.59, 2.62] pp (two simultaneous intervals). Four covariance intervals span zero. Only four-presentation held-out THINGS Ridge beats its null (+0.95 pp [0.01, 1.94] in 20 comparisons).

**Frozen rankings across datasets.** Use domestic Ridge and MLP for seven nonpilot held-out participants per dataset. Both conditions share 159 development concepts, 738 queries and 1,573 references. The half quota keeps 3,172 of 6,403 directed pairs per person. Signed support (Semantic agreement in main Table 5) orients the semantic margin by the spatial decision. Agreement puts matching signs first, then uses absolute spatial margin. Absolute uses that margin alone. People and concepts have equal weight.

| | | Signed support gain, single presentation | | Change in gain, four minus one | |
|---|---|---|---|---|---|
| Dataset | Readout | vs. agreement | vs. absolute | vs. agreement | vs. absolute |
| THINGS-EEG2 | Ridge | $+0.28$ $[-1.03, +1.50]$ | $+3.75$ $[+0.78, +6.95]$ | $+0.69$ $[-0.81, +2.25]$ | $+0.82$ $[-2.08, +3.73]$ |
| THINGS-EEG2 | MLP | $+0.67$ $[-0.76, +2.01]$ | $+4.32$ $[+1.46, +7.43]$ | $+0.16$ $[-1.39, +1.78]$ | $+0.73$ $[-2.62, +4.13]$ |
| Alljoined-1.6M | Ridge | $+0.25$ $[-1.14, +1.58]$ | $+1.04$ $[-2.04, +3.96]$ | $-0.30$ $[-2.13, +1.52]$ | $-0.03$ $[-3.32, +3.32]$ |
| Alljoined-1.6M | MLP | $+0.04$ $[-1.23, +1.21]$ | $-0.44$ $[-3.09, +2.64]$ | $+0.11$ $[-1.55, +1.72]$ | $+1.08$ $[-2.57, +5.18]$ |

Across three quotas (1,556, 3,172 and 4,756), the six single-presentation THINGS gains over absolute margin exclude zero. The other single-presentation and 24 four-minus-one intervals span zero (48 simultaneous 95% intervals in pp). Four-presentation semantics with fixed spatial decisions yields 16 half-coverage intervals spanning zero.

**Donor participant and image controls.** We hold spatial decisions fixed. Six other people in the same dataset view either the same image or all nonzero within-concept query shifts. Intervals resample concepts with people fixed.

| Dataset | Readout | Own − same signed | Own − same agreement | Same − other signed |
|---|---|---|---|---|
| THINGS-EEG2 | Ridge | $-0.07$ $[-0.48, +0.32]$ | $-0.26$ $[-0.71, +0.18]$ | $+4.94$ $[3.47, 6.52]$ |
| THINGS-EEG2 | MLP | $-0.18$ $[-0.59, +0.22]$ | $-0.30$ $[-0.74, +0.12]$ | $+5.82$ $[4.06, 7.68]$ |
| Alljoined-1.6M | Ridge | $-0.19$ $[-0.71, +0.33]$ | $-0.08$ $[-0.56, +0.40]$ | $+1.95$ $[0.41, 3.49]$ |
| Alljoined-1.6M | MLP | $-0.10$ $[-0.53, +0.33]$ | $-0.16$ $[-0.66, +0.34]$ | $+1.71$ $[0.10, 3.29]$ |

Same-image donors beat CLIP and $R_8$ candidate distances by 4.71–5.55 pp in THINGS-EEG2. All four Alljoined intervals include zero.

**Cross-dataset donors.** Same-image donors beat other-image or spatial-only in 4/8 groups each. Own beats donors in 1/8; donors beat own in 0/8. Source effects favor THINGS in 3/4 (simultaneous intervals).

**Fixed thresholds with a single presentation.** Each signed-support cutoff is the source median from 7,006 training images and two people. Both tests share the cutoff and gallery. People and concepts have equal weight. Accuracy divides retained correct mass by retained mass.

| | | Domestic (%) | | Foreign (%) | | Foreign − domestic (pp) | |
|---|---|---|---|---|---|---|---|
| Training dataset | Readout | Coverage | Accuracy | Coverage | Accuracy | Coverage | Retained accuracy |
| THINGS-EEG2 | Ridge | 48.84 | 55.29 | 48.21 | 52.48 | $-0.63$ $[-2.68, +1.55]$ | $-2.82$ $[-7.68, +1.68]$ |
| THINGS-EEG2 | MLP | 46.02 | 56.16 | 45.52 | 52.75 | $-0.50$ $[-2.57, +1.72]$ | $-3.41$ $[-8.22, +1.30]$ |
| Alljoined-1.6M | Ridge | 48.08 | 52.41 | 48.84 | 51.04 | $+0.76$ $[-1.28, +2.80]$ | $-1.37$ $[-5.87, +3.19]$ |
| Alljoined-1.6M | MLP | 45.33 | 51.56 | 45.99 | 50.59 | $+0.65$ $[-1.41, +2.95]$ | $-0.97$ $[-5.84, +3.80]$ |

All eight transfer intervals include zero. Gains over full coverage clear zero only within THINGS-EEG2: Ridge $+4.47$ $[2.24, 6.95]$ and MLP $+5.14$ $[2.82, 7.58]$ pp.

#### 10. Training windows, evaluation panels and secondary controls

**EEG and frozen representations.** Inputs are 63 channels and 250 samples at 250 Hz. Sub-01/02 baseline correction precedes antialiasing and cropping. Training presentations set whitening and normalization before averaging. Shared and semantic features have 1,440 and 1,024 dimensions. Linear and 256-wide GELU probes predict 16 $D_4$, 64 $R_8$ and 256 $R_{16}$ coordinates. Transfer uses the source training center.

**Separate visual-input geometry intervention.** Visual inputs (128 images, 32 concepts, three seeds). Own vs. donor tests full-depth $D_4$ and fixed-$D_4$ $R_8$. Same-image cosine gains average three same-concept alternatives (eight simultaneous 95% intervals).

| Output depth readout | Output recurrence across seeds | Alignment with injected direction |
|---|---|---|
| $D_4$ (DA3) | $+0.032$ $[-0.028, +0.093]$ | $+0.218$ $[+0.138, +0.289]$ |
| $D_4$ (DAv2) | $+0.094$ $[+0.032, +0.153]$ | $+0.220$ $[+0.140, +0.297]$ |
| $R_8$ (DA3) | $+0.009$ $[-0.025, +0.044]$ | $+0.039$ $[+0.006, +0.072]$ |
| $R_8$ (DAv2) | $+0.008$ $[-0.030, +0.045]$ | $+0.032$ $[-0.010, +0.071]$ |

All 16 win-rate intervals include zero. The eight-case replay matches all 16 RGB outputs and finds a pre-RGB location decline in six cases.

**Appearance-neighbor geometry.** Rank candidates by four equally weighted RGB distances (luminance, contrast, color, edges), scaled on 2,160 training pairs. Nearest and farthest strata each have 4,890 comparisons across 163 concepts, excluding the cyclic donor.

| | Matched gain on nearest candidates | | Gain change, nearest $-$ farthest | |
|---|---|---|---|---|
| NICE input target | Over cyclic | Over coarse | Over cyclic | Over coarse |
| Projected CLIP | $+9.06$ $[+5.26, +12.60]$ | $+6.32$ $[+3.63, +8.92]$ | $-1.51$ $[-4.44, +1.35]$ | $+1.15$ $[-2.19, +4.54]$ |
| Block 16 | $+9.20$ $[+5.84, +12.52]$ | $+6.61$ $[+4.17, +9.05]$ | $-3.66$ $[-6.73, -0.57]$ | $-0.65$ $[-3.79, +2.64]$ |
| Block 32 | $+8.26$ $[+5.15, +11.52]$ | $+6.05$ $[+3.61, +8.57]$ | $-2.39$ $[-5.05, +0.35]$ | $-0.37$ $[-3.84, +3.09]$ |

Simultaneous 95% intervals (pp) cover 12 contrasts. Block 16 loses cyclic gain on closer pairs (sub-08 development, monocular depth).

**Joint holdout eligibility.** Each new participant requires 24 new concepts with two EEG-paired images each. Removing castle leaves all 16 first-panel NOD intervals spanning zero.

| Dataset | Count under the current screen | Remaining evidence |
|---|---|---|
| NOD | 0 unused of 148 screened concepts | New concepts |
| Neuro-3D | At most 19 nonmatching source labels | Static-frame onset and whitening provenance |
| Alljoined1 | At most 21 repeated foreground labels | Finer labels in the shown crops |
| FOODEEG | At most 27 pairs (91 images, 37 text overlaps) | Object labels, image provenance and use terms |
| Greene & Hansen | At most 29, or 26 if the related roster matches | EEG labels, stimuli, trial and sensor IDs |
| Yang et al. | 83 of 181 pairs pass text and nonempty-file screens | Electrode order and image checks |
| Chen et al. | 320 single-image food stimuli | Single-image EEG and onsets |

**Where can TRAIN and DEV windows share EEG samples?** We joined archived sub-08 trigger onsets to fixed TRAIN and DEV image identities. A DEV window overlaps if it shares at least one sample with a TRAIN candidate in the same recording segment. The table counts overlaps among eligible DEV windows.

| Window (ms from onset) | Archive DEV candidates | Four presentations in later benchmark |
|---|---|---|
| $[0, 160)$ | 0/6,577 | 0/6,520 |
| $[0, 1000)$ | 6,576/6,577 | 6,519/6,520 |
| $[-200, 1000)$ | 6,577/6,577 | 6,520/6,520 |

**What the archive establishes.** TRAIN and DEV have disjoint image and concept identities. Project channel scaling used four slots for each of 13,910 TRAIN images. If both historical four-slot sets came from this archive, at least 6,462 of 6,520 nominal DEV $[0, 1000)$ ms windows share TRAIN samples for any slot choice. TRAIN images with exactly four archive events already establish the bound. No candidate windows overlap at $[0, 160)$ ms.

**What remains unresolved.** Historical slot identities, filter support and upstream whitening membership are unverified. The short-window refit in Sec. 11 uses raw-derived windows. Later cross-person probes for sub-08 use six other people for fitting.

**Reading main Sec. 5.1.** On the exposed historical 100-concept sub-08 panel (999 valid images, 4,491 pairs), fixed predictions yield 52.10% squared distance and 58.20% centered cosine single-query accuracy, a $+6.10$ pp change $[4.43, 7.78]$. This is a scoring comparison. On the 163-concept DEV panel, the historical heads' matched-minus-bilinear $R_8$ gains are $+5.60/+3.14/+1.47$ pp for sub-08/01/02. Main Table 3 and Fig. 4 use refitted probes on the DEV panel. Their MLP matched-minus-coarse gains are $+7.79/+3.59/+2.57$ pp. The historical 100-concept DA3 exchange figures in the main text use joint assignment.

**Two frozen CNNs on 100 held-out concepts.** On 8,982/9,000 directions with shared support, cosine centered on 624 fitting concepts raises accuracy from 50.93/51.01% to 54.56/53.83%. Gains are $+3.63$ $[1.79, 5.54]$ and $+2.81$ $[0.93, 4.77]$ pp (four simultaneous intervals).

#### 11. Recovery and reliability with entire sessions held out

**Historical session 4 comparison.** Fit whitening, normalization and a fresh encoder on sessions 1–3. Compare the same four images per concept across a source session and session 4, averaging two presentations per image. Historical DEV has 76 images in 19 concepts (sub-01) and 56 images in 14 concepts (sub-02).

**Single-query comparison.** Score 228/168 directed comparisons with DAv2-Small, a TRAIN-only center and common residual cells. Controls use the next same-concept image in a fixed order or interpolate the query's $D_4$.

| Participant | Control | Source gain (pp) | Session 4 gain (pp) | Session 4 − source (pp) |
|---|---|---|---|---|
| sub-01 | Cyclic | $+7.46$ $[-2.63, +17.98]$ | $+7.46$ $[-6.14, +18.86]$ | $+0.00$ $[-19.74, +15.79]$ |
| sub-01 | Coarse | $+3.07$ $[-4.39, +10.96]$ | $+2.63$ $[-5.26, +9.65]$ | $-0.44$ $[-9.65, +9.21]$ |
| sub-02 | Cyclic | $+10.12$ $[-1.79, +22.02]$ | $+1.79$ $[-5.95, +10.12]$ | $-8.33$ $[-20.83, +3.57]$ |
| sub-02 | Coarse | $+7.14$ $[-1.79, +14.88]$ | $+0.60$ $[-6.55, +8.33]$ | $-6.55$ $[-16.67, +4.76]$ |

Gains are matched minus control. All twelve simultaneous 95% intervals include zero. The paired interval uses the same concept resamples for both sessions.

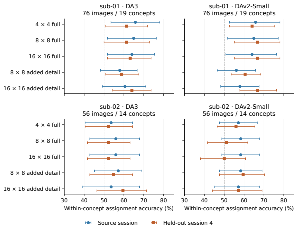


**Historical joint assignment on the same images.** Points and marginal 95% intervals use pairs of EEG queries. The $R_8$ DA3 estimate on session 4 is 58.77% [50.88, 67.54] for sub-01 and 54.76% [42.86, 64.29] for sub-02. These values describe the joint task. The table above tests single-query gains over within-concept controls.

**Short windows with shared sessions (sub-08).** TRAIN and DEV can share sessions here. Average eight 20 ms bins per channel over 0–160 ms (63 channels, 1 kHz). Fit scaling and Ridge ($\lambda = 0.1n$) on 13,883 TRAIN images with DA3 targets and no added baseline or whitening. Across 14,670 directions, matched/cyclic/coarse scores are 54.70/49.02/50.67%. Gains are $+5.68$ $[3.18, 8.12]$ and $+4.02$ $[2.36, 5.67]$ pp (simultaneous 95% concept intervals).

**Recovery before and after onset in held-out sessions.** Fit Ridge and scaling separately for $[-160, 0)$ and $[0, 160)$ ms, using TRAIN from the other three sessions. Both windows use the same events and two presentations per held-out image. All 6,520 DEV slots per person pass both boundary checks, and all post-onset scores replay exactly. Average each concept's two held-out folds before resampling 163 DEV concepts per person.

| Participant | Control | Pre gain (pp) | Post gain (pp) | Post − pre gain (pp) |
|---|---|---|---|---|
| sub-08 | Cyclic | $+0.93$ $[-1.31, +3.16]$ | $+3.46$ $[1.04, 5.88]$ | $+2.53$ $[-0.62, +5.77]$ |
| sub-08 | Coarse | $+0.39$ $[-1.07, +1.87]$ | $+2.62$ $[1.00, 4.17]$ | $+2.23$ $[0.38, 4.12]$ |
| sub-01 | Cyclic | $-0.41$ $[-2.92, +2.07]$ | $+3.74$ $[1.32, 6.16]$ | $+4.15$ $[0.91, 7.36]$ |
| sub-01 | Coarse | $-0.40$ $[-2.04, +1.22]$ | $+3.20$ $[1.79, 4.61]$ | $+3.60$ $[1.89, 5.29]$ |
| sub-02 | Cyclic | $-0.82$ $[-3.08, +1.43]$ | $+3.26$ $[0.83, 5.65]$ | $+4.08$ $[1.69, 6.49]$ |
| sub-02 | Coarse | $-0.13$ $[-1.61, +1.35]$ | $+2.66$ $[1.12, 4.14]$ | $+2.79$ $[1.29, 4.24]$ |

Gains are matched minus control (18 simultaneous concept intervals). Every post-onset gain and five post-minus-pre differences exclude zero. All pre-onset gain intervals include zero. These early windows can include activity from the preceding image or sequence effects. Counts by fold (sessions 1–4) are 79/84/82/81, 85/78/84/79 and 83/80/75/88 concepts for sub-08/01/02. The earlier 180 ms joint-assignment pilot (48 pairs per person) gave $+4.17$ pp post minus pre for sub-01/02, with both intervals spanning zero.

**Expanded reliability rankings at the 45/90 quota (sub-08).** Keep 45/90 fixed post-onset decisions per concept. Fit error models to concept-cross-fitted TRAIN residuals. Directional, isotropic and diagonal scores retain 55.08/55.25/55.09% correct. Plane margin and cosine gap retain 55.22/55.39%, signed support and absolute semantic gap 54.46/53.28%, and cyclic and candidate geometry 52.43/53.46%. Directional minus isotropic is $-.17$ $[-.74, +.40]$ pp, directional minus diagonal is $-.01$ $[-.52, +.50]$, and signed minus directional is $-.62$ $[-1.85, +.59]$. The intervals retain the eight-comparison adjustment ($n = 163$ concepts). An optimizer warning leaves the learned head and its contrast unreported.

#### 12. What does the Fig. 5 depth metric detect?

**Fixed-teacher replay on cached RGB.** With identical sub-08 RGB, the original run matched 384/384 DAv2-Small maps and 275/384 DA3 maps. Two seeded DA3 passes on 384 cached outputs reproduce depth, masks and 1,152 judgments, across the fixed output archive. The archived CPU state replays the unseeded sky branch. The table describes the historical 32-stimulus DA3 analysis (eight concepts, two probe seeds, one generation seed, two semantic settings, four simultaneous 95% intervals). Fig. 5 uses DAv2 below.

| Semantic readout | Geometry control | Input advantage (pp) | Output advantage (pp) | Input − output (pp) |
|---|---|---|---|---|
| Original | Cyclic donor | 17.71 | 0.00 | 17.71 [0.00, 37.50] |
| Original | Zero detail | 9.90 | −1.04 | 10.94 [−6.77, 27.08] |
| Linear | Cyclic donor | 17.71 | −1.04 | 18.75 [0.52, 38.54] |
| Linear | Zero detail | 9.90 | −2.08 | 11.98 [−6.77, 29.17] |

**Where do changes appear during denoising?** All 16 outputs replay with layout, semantics and noise fixed. The trace relates changes in the $8^2$ input to latent block RMS and RGB mean absolute changes. It subtracts the mean of three rotated correlations.

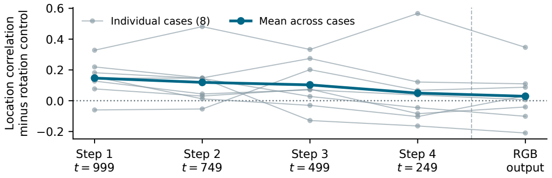


Location correspondence falls in six of eight cases (exact sign $p = .289$), by $-.097$ on average (case-bootstrap 95% interval $[-.214, +.032]$). The trace tests location correspondence. The stage of loss remains unresolved.

**Calibration with known flips at each scoring scale.**

| Grid | True direction energy | DAv2-Small vs. control | DA3 vs. control |
|---|---|---|---|
| 16 × 16 | 4.79% | −.0034 [−.0101, +.0018] | −.0022 [−.0070, +.0037] |
| 32 × 32 | 11.34% | −.0013 [−.0076, +.0048] | −.0009 [−.0123, +.0136] |
| 64 × 64 | 25.52% | +.0093 [+.0005, +.0181] | +.0214 [−.0012, +.0506] |
| 128 × 128 | 53.01% | +.0300 [+.0129, +.0485] | +.0703 [+.0215, +.1325] |
| 500 × 500 | 100.00% | +.0501 [+.0243, +.0779] | +.1224 [+.0463, +.2227] |

Energy is median squared-norm retention. Both contrasts pass from $64^2$ for DAv2-Small and $128^2$ for DA3. At $500^2$, the teachers order 20/24 and 22/24 pairs correctly, respectively, and each resolves both states in 4/24.

**Fig. 5 joint score on known flips.** On 24 controlled flips, DAv2 joint assignments are 13/24 at the original $8^2$ score grid and 22/24 at $128^2$. Gains over chance are $+4.17$ $[-20.83, 29.17]$ and $+41.67$ $[25.00, 50.00]$ pp (six simultaneous intervals). Independent decisions resolve both states in 0/24 and 5/24. The $8^2$ direction and specificity intervals both include zero.

**Main Fig. 5 scored at the same scale before and after generation.** Sub-08 depth and semantics come from EEG. Exchange swaps detail within a concept at fixed recipient layout and semantics. The panel has 128 DAv2-Small references and 768 outputs (32 concepts, 192 pairs, three seeds). The generator uses $16^2$ inputs. Scoring expands them by nearest neighbors and projects maps where indicated. Sec. 10 reports a separate visual-input test.

**Independent state choices.** For a pair to count, each query must be closer to its own recipient reference. At $64^2$ and $128^2$, both inputs are correct in 9/192 Own and 4/192 Exchange pairs. Both outputs are correct in 71/576 pairs per arm at $64^2$ and 68/576 at $128^2$. Output-minus-input gain intervals include zero. Visibility in natural RGB outputs remains unmeasured.

| Score grid | Input gain (pp) | Output gain (pp) | Output − input (pp) |
|---|---|---|---|
| $8^2$ (historical Fig. 5, exact replay) | +25.52 | +0.17 | −25.35 [−38.19, −12.50] |
| $128^2$ (full replay) | +25.52 | +0.69 | −24.83 [−41.90, −6.71] |
| $64^2$ (common projection) | +25.52 | +0.87 | −24.65 [−41.67, −6.60] |
| $16^2$ (common projection) | +25.52 | +0.35 | −25.17 [−40.97, −8.68] |

**Joint assignment.** The table compares both possible assignments of each pair and reports the Own minus Exchange gain. The $8^2$ interval is marginal. The $16^2$ interval adjusts for six contrasts and the $64^2/128^2$ intervals for twelve. Known flips pass directional and specificity checks under the same $64^2/128^2$ projections.

**Equal-dose visual and EEG residuals.** At matched $R_8$ RMS on 32 images, DA3 input/output gains are $+77.08/+0.69$ pp (visual) and $+37.50/0.00$ pp (EEG). DAv2-Small $128^2$ output gains are $-1.39$ $[-5.56, 0.00]$ pp (visual) and $+0.69$ $[-2.08, +3.47]$ pp (EEG), with eight adjusted intervals. Fixed midpoints and $[0, 1]$ bounds permit $2/4/8\times$ contrast for 16/2/0 images in both sources, with zero complete concepts at $2\times$.

#### 13. How scoring and EEG averaging affect the results

**Spatial coordinates.** Each image has a relative depth map. Align depth signs so larger values mean nearer, clip and rescale within image using the valid 5th and 95th percentiles, and pool on shared valid support. $D_4$ is the $4 \times 4$ layout. $R_8 = D_8 - U(D_4)$ and $R_{16} = D_{16} - U(D_8)$ are the added details under nearest-neighbor expansion $U$. For a prediction or candidate residual $v$ on the common valid cells, compare inner products of

$$z(v) = \frac{(v-\mu)/s}{\|(v-\mu)/s\|_2}.$$

The same template $\mu$, scale $s$, and support apply to query and both candidates. Unscaled cosine uses $s = 1$. Zero norms are recorded as undecided and count half; no zero norms occur in this reanalysis.

**Sensitivity of the original scoring comparison.** Main Table 2 fixes the historical sub-08 predictor, DA3 candidates and all 14,670 directed comparisons on 163 development concepts. The DAv2-Small template uses 13,909 valid TRAIN images, and the DA3 template uses 11,095 in the probe's fitting split. Both the teacher and fit membership differ. Main Table 3 averages two refit seeds per readout (linear 61.59%, MLP 61.23%). Main Table 5 and Fig. 6 fix the seed-42 MLP (61.22% before ranking).

| Score | Correct (%) | Prespecified comparison (pp, simultaneous 95% CI) |
|---|---|---|
| Squared distance | 52.73 | |
| Raw cosine | 55.31 | |
| DAv2 TRAIN-centered cosine | 59.36 | |
| DA3 TRAIN-centered cosine | 58.28 | vs. DAv2 center: $-1.08$ $[-1.77, -0.40]$ |
| DA3 center and standard deviation | 58.14 | vs. unscaled DA3: $-0.14$ $[-0.63, +0.38]$ |
| | | vs. DAv2 center: $-1.22$ $[-1.99, -0.47]$ |

**Single presentations and their average.** Both readouts retain a matched-minus-cyclic advantage on single presentations. Scores are higher with averaged EEG. The two TRAIN-fitted readouts use DA3 and the same 1,630 images in 163 concepts. We average the four presentation-level scores and the two fitting seeds before concept-level resampling. Historical session assignments for the four presentations are unresolved.

| Readout | Single (%) | Four (%) | Single matched − cyclic (pp) | Four − single (pp) |
|---|---|---|---|---|
| Linear | 56.50 | 61.59 | 6.89 $[5.10, 8.64]$ | 5.09 $[3.85, 6.33]$ |
| MLP | 56.71 | 61.23 | 6.78 $[5.10, 8.42]$ | 4.51 $[3.48, 5.53]$ |

**EEG-only fidelity prediction and post-generation selection (main Fig. 2).** For prediction, fixed sub-08 pre-prior embeddings yield four log-error risks ($D_4$, $R_8$, $R_{16}$ and semantics). Risk fitting excludes both the 32 quality-fitting and 32 exposed evaluation concepts. Heads use only EEG-derived inputs. Offline credits use three seeds and three alternatives per image. Brier uses 128 images, with 32 reserved for calibration and 96 for calibrated Brier and 50%-retained credit.

| | | Semantic target | | | $R_8$ target | |
|---|---|---|---|---|---|---|
| Predictor | Brier | Cal. Brier | Credit | Brier | Cal. Brier | Credit |
| Constant | .23906 | .23832 | .57870 | .25342 | .25053 | .51852 |
| Full linear | .27224 | .23975 | .60995 | .27605 | .25244 | .48380 |
| Full MLP | .25355 | .24475 | .55903 | .25470 | .25293 | .49421 |
| PCA4 | .24353 | .23977 | .57523 | .25248 | .25011 | .50000 |
| Random projection4 | .24654 | .23814 | .61806 | .25487 | .25514 | .48727 |
| Spatial risks | .24278 | .23892 | .60301 | .25190 | .25184 | .50926 |
| Four risks | .24001 | .23935 | .59838 | .25255 | .25048 | .50579 |
| Swapped risks | .23947 | .24083 | .56944 | .25313 | .25061 | .52199 |
| Full + risks | .25448 | .24102 | .59722 | .25199 | .25096 | .52315 |

Four risks reduce semantic Brier relative to the full MLP by .01354 (adjusted 95% interval [.00582, .02122]). Differences relative to PCA4, random projection4 and swapped risks are uncertain. The six $R_8$ prediction contrasts and five retention contrasts per target all span zero.

**Post-generation selection.** Choose among three cached outputs per image for 128 images. TRAIN-fitted own $R_8$ earns 55.47% spatial credit vs. 55.73% uniform expectation (adjusted difference $-0.26$ pp, $[-8.33, +7.03]$), 53.12% for new semantic Linear and 60.94% each for cyclic $R_8$ and old post-prior semantic selection. This comparison reads the generated images at scoring time.

**Why depth residuals.** Subtracting the coarser field gives an explicit interpolation control. RGB edges resolve known boundaries, but natural-edge matching yielded 66 image pairs in only nine concepts with at least two pairs (30 required). Finer $R_{16}$ transfer remains unresolved (Sec. 1).

#### 14. Does the current presentation help rank decisions?

**TrustScore-style gallery baseline.** Assign eight other centered unit-$R_8$ references to the nearer candidate, excluding ties and including each candidate in its own neighborhood. Rank by the log ratio of nearest rejected to chosen distances [14]. Keep 45/90 decisions per concept without correctness labels.

| | Retained accuracy (%) | | | Neighbor ratio minus comparison (pp) | |
|---|---|---|---|---|---|
| Participant | Abs. margin | Agreement | Neighbor | Abs. margin | Agreement |
| sub-08 | 66.79 | 83.65 | 63.04 | $-3.75$ [$-5.44, -2.03$] | $-20.61$ [$-23.00, -18.24$] |
| sub-01 | 58.98 | 66.63 | 57.40 | $-1.58$ [$-3.34, +0.15$] | $-9.23$ [$-11.48, -6.98$] |
| sub-02 | 55.13 | 63.31 | 53.69 | $-1.45$ [$-3.30, +0.33$] | $-9.63$ [$-11.95, -7.30$] |

Six simultaneous intervals. The 1,571-feature head hit the 1,000-iteration cap in sub-01 fold 2 and sub-02 fold 1. Analytic baselines use all five folds.

**Candidate separation without query EEG.** Candidate cosine distances retain 63.48/56.44/54.36% ($R_8$) and 62.09/56.03/53.35% (CLIP) on sub-08/01/02. All six differences from semantic support are below zero (12 simultaneous contrasts).

**Same-image donors at the main 45/90 quota.** On sub-08's fixed decisions, Own retains 83.65% and the mean of sub-01/02 donors retains 71.21% ($+12.45$ [10.80, 14.10] pp). Own minus the two-donor mean is $-6.22$ [$-7.89, -4.55$] pp in sub-01 and $-8.00$ [$-9.67, -6.30$] pp in sub-02 (nine simultaneous 95% intervals). These swaps change both participant and readout.

**Ranking within difficulty strata.** Stratify each concept by donor support and $R_8$ distance (three bins each); keep 5/10 per bin. Own minus mean donor is $+14.37$ [12.98, 15.81], $-1.31$ [$-2.76, +0.17$], and $-3.03$ [$-4.56, -1.51$] pp for sub-08/01/02. Own beats within-bin hash (six simultaneous intervals). Native semantic accuracies differ (75.84/61.75/60.47%); bins may retain difficulty differences.

**Two-feature correctness model.** Fit and scale L2 logistic models on margin and signed support in four concept folds. Rank 45/90 fixed queries per concept in the fifth. All 15 fits converged in 6–8 iterations.

| Participant | Retained accuracy (%) | Minus abs. margin (pp) | Minus agreement (pp) |
|---|---|---|---|
| sub-08 | 84.17 | $+17.38$ [15.39, 19.33] | $+0.52$ [$-0.02, +1.04$] |
| sub-01 | 67.09 | $+8.11$ [6.15, 10.02] | $+0.46$ [$-0.26, +1.19$] |
| sub-02 | 63.37 | $+8.23$ [6.19, 10.24] | $+0.05$ [$-0.53, +0.63$] |

The six simultaneous intervals support gains over absolute margin. The incremental gains over agreement remain uncertain on this development panel.

**Alternative semantic readouts.** For sub-08, NICE, UBP, CLIP Ridge and SwAV Ridge retain 80.20/80.16/77.53/81.83%. Matched-null gains of $+1.77/+2.00/+1.41/+2.20$ pp have positive adjusted intervals. Shared image difficulty remains possible.

**Same person and image, different EEG presentation.** On 1,630 images with nine alternatives per image, retain 2/4 fixed seed-42 $R_8$ decisions using either the current presentation or each of three other presentations of the same image. Average donor accuracies. Use a TRAIN-only center, fixed hash ties and nine simultaneous 95% concept intervals.

| Participant | Same-slot (%) | Donor-slot (%) | Same minus donor (pp) |
|---|---|---|---|
| sub-01 | 58.80 | 58.87 | $-0.07$ [$-0.39, +0.25$] |
| sub-02 | 57.66 | 57.77 | $-0.11$ [$-0.42, +0.20$] |
| sub-08 | 68.06 | 68.19 | $-0.13$ [$-0.34, +0.09$] |

Own-slot gains over spatial margin are $+4.08/+4.82/+9.33$ pp and over hash $+5.01/+5.15/+11.40$ pp (all six adjusted intervals above zero). Event identity is known in sub-01/02; historical sub-08 slots lack that binding.

**Donors across two held-out sessions.** For sub-01/02, Ridge fits TRAIN from two other sessions (0–160 ms), keeping 2/4 fixed decisions per pair. Own minus cross-session donor (swap average) is $-.06$ [$-.19, +.06$] and $+.03$ [$-.08, +.15$] pp; minus within-session donor $-.05$ [$-.19, +.10$] and $+.05$ [$-.06, +.17$] pp; minus hash $+.68$ [$-.01, 1.36$] and $+.88$ [$.17, 1.58$] pp (six simultaneous intervals). Single-trial semantics score 51.14/50.87%.

**Two versus four presentations with the same held-out sessions (sub-08).** Averaging all four presentations raises signed retention from 54.07% to 55.43% ($+1.36$ [0.25, 2.45] pp) relative to the mean of two complementary cross-session pairs. Both use the same geometry/CLIP Ridge fit on the other two sessions (0–160 ms) and keep 45/90 decisions per concept. With four presentations, signed minus hash is $+1.81$ [0.61, 3.07] pp. Differences from plane margin (55.45%) and binary agreement (55.08%) include zero (ten simultaneous intervals).

#### 15. Scoring equations and reproducibility

**Fixed candidate decision.** Main Eqs. 5–7 use centered unit vectors $a, b, q$ and $c = (a + b)/2$, $\delta = \|a - b\|_2$, $u = (a - b)/\delta$, $m = u^\top(q - c)$. The squared-distance margin is $H(q) = \|q - b\|^2 - \|q - a\|^2 = 2\delta m$. On two observations, correct-minus-swapped joint assignment is $H(q_A) - H(q_B)$, so a common additive candidate preference cancels. The two single-query decisions still depend on its sign and magnitude. With zero residual mean and isotropic variance, the directional score is proportional to the absolute centered-cosine gap $\delta|m|$.

**Scores of the same decision.** For each fitting image, use one normalized residual relative to its own reference. Let their mean and sample covariance be $\mu, S$. Directional covariance uses $\Sigma = (S + \operatorname{diag} S)/2 + \epsilon I$, $\epsilon = \max(10^{-6}\operatorname{mean}(\operatorname{diag} S), 10^{-12})$, and ranks by $\operatorname{sign}(m)(m - u^\top\mu)\delta/(u^\top\Sigma u)$. Isotropic uses $\operatorname{tr}(\Sigma)I/64$ with the same mean; diagonal uses $\operatorname{diag}\Sigma$. Other rankings are $|m|$, $\delta|m|$, candidate-only $\delta/\sqrt{u^\top\Sigma u}$, semantic gap magnitude, signed semantic support, the directional score with cyclic other-image evidence and the learned correctness head. The semantic gap uses pre-prior CLIP cosine differences. The head uses shared EEG features (1,440), signed direction and midpoint (64 each), $|m|$, $\delta$ and signed semantic support (1,571 total), with fitting-only scaling, fixed L2 regularization and a 1,000-iteration cap.

**Folds and inference.** Sort 163 concepts by SHA-256 and assign round-robin to five folds. Estimate statistics on four folds without refitting the recovery probe. Rank the held-out fold's 90 directed queries per concept and retain 45. Break confidence ties by fixed anonymous query and candidate hashes. Keep half credit for unresolved base decisions. Primary intervals use 20,000 paired whole-concept draws, seed 20261117, family-four quantiles 0.00625 and 0.99375. They condition on the fitted models and retained sets. Equal-capacity nested heads keep semantic support, cosine gap and candidate distance in all arms. Changing the fourth input from directional covariance to ordinary margin, isotropic, cyclic or candidate geometry did not establish a directional benefit in the main comparison.

**Covariance sensitivity.** Reuse each fold's $S, \mu$ and fixed decisions. Give off-diagonals weights $0, .25, .5, 1$ and use projected variance or standard deviation. Relative to the same-exponent isotropic baseline, differences span $-0.76$ to $+0.40$ pp across three development participants. All 24 simultaneous 95% concept-bootstrap intervals include zero (20,000 draws, seed 20261124).

**Generation and scope.** Record stimulus and donor IDs, checkpoints, seeds, supported score cells and failures. Replay the same RGB before estimating sensitivity at the observed amplitude. The declared depth endpoint measures relative attributes; known-flip calibration and natural-output fidelity require separate evidence. AttrEcho [27], ProtoEEG [29] and SGDM [19] address related reconstruction questions on their own tasks. Main Fig. 1a shows one of two fixed EEGRFusion training examples (sub-08, four presentations averaged). Among ten dog images, the viewed stimulus ranked first at the EEG encoder for both cases and fourth and ninth after the flow mapping, respectively. The second output depicted a bear-like animal. Panel b uses a separate, frozen 32-image development panel. The two silicone stimuli were scored against each other using training-centered, unit-normalized $R_8$ coordinates.
